# Single-nucleus RNA-seq reveals that MBD5, MBD6, and SILENZIO maintain silencing during epigenetic reprogramming in pollen

**DOI:** 10.1101/2022.09.29.510154

**Authors:** Lucia Ichino, Colette L. Picard, Jaewon Yun, Meera Chotai, Shuya Wang, Evan Kai Lin, Ranjith K. Papareddy, Yan Xue, Steven E. Jacobsen

## Abstract

Silencing of transposable elements (TEs) drove the evolution of numerous redundant mechanisms of transcriptional regulation. Arabidopsis MBD5, MBD6, and SILENZIO act as TE repressors downstream of DNA methylation. Here we show via single-nucleus RNA-seq of developing male gametophytes that these repressors are critical for TE silencing in the pollen vegetative cell, which undergoes epigenetic reprogramming causing chromatin decompaction to support fertilization by sperm cells. Instead, other silencing mutants (*met1*, *ddm1*, *mom1*, *morc*) show loss of silencing in all pollen nucleus types and somatic cells. We found that TEs repressed by MBD5/6 gain accessibility in wild-type vegetative nuclei despite remaining silent, suggesting that loss of DNA compaction makes them sensitive to loss of MBD5/6. Consistently, crossing *mbd5/6* to *histone 1* mutants, which have decondensed chromatin in leaves, reveals derepression of MBD5/6-dependent TEs in leaves. MBD5/6 and SILENZIO thus act as a silencing system especially important when chromatin compaction is compromised.

## INTRODUCTION

Eukaryotes have evolved numerous molecular strategies to establish and maintain silencing of genes and repetitive elements. Epigenetic modifications of histones and DNA work jointly with chromatin remodelers and histone variants to make DNA inaccessible to the transcriptional machinery (Beisel and Paro, 2011). During development, some cells undergo a process of global epigenetic reprogramming, which entails the erasure of different layers of repression (Feng et al., 2010). An example of this occurs in the mammalian primordial germ cells (PGCs), which undergo a genome-wide loss of DNA methylation to reestablish genomic imprints (Hajkova, 2011). This transient relaxation of epigenetic repression creates a window of opportunity for mobile elements to reactivate and transpose, with potentially harmful consequences to genome integrity (Zamudio and Bourc’His, 2010).

In the flowering plant *Arabidopsis thaliana*, germline epigenetic reprogramming involves some changes in histone variants, histone tail marks, and non-CG methylation, but CG methylation levels are not globally erased like in mammals, thus allowing for transgenerational epigenetic inheritance of DNA methylation epialleles (Gehring, 2019; Kawashima and Berger, 2014). On the other hand, extensive rewiring of chromatin structure and partial loss of CG methylation occur in reproductive “companion cells”, which function as supporting cells during fertilization and embryogenesis (Kawashima and Berger, 2014). The development of male gametophytes in Arabidopsis starts with a meiotic event that generates four identical haploid microspores from a Microspore Mother Cell. Each microspore subsequently undergoes an asymmetric mitotic division that produces a Vegetative Cell (VC) and a Generative Cell (GC). The GC is engulfed within the cytoplasm of the VC, thus creating the bicellular pollen grain. The GC then undergoes another round of mitosis that produces two identical Sperm Cells (SCs), forming the tricellular pollen grain. The SCs constitute the germline, while the VC is a supporting cell that is responsible for pollen tube growth, which allows delivery of the SCs to the female gametes for fertilization (Johnson et al., 2019). The generation of pollen grains, from microspores to the tricellular stage, occurs over a time frame of about three days (Alvarez-Buylla et al., 2010; Sanders et al., 1999).

The VC and the SC undergo remarkably different chromatin reorganization events that eventually lead to the formation of two small and highly compacted sperm nuclei (SN) and a very large and decondensed vegetative nucleus (VN). This striking difference is apparent in microscopy images of DAPI stained mature pollen (Borges et al., 2012). More specifically, chromatin decompaction in the VN occurs, at least in part, because of the depletion of the linker histone H1, which begins at the late microspore stage (He et al., 2019) and because of the active removal of the centromere-specific histone H3 variant (CenH3) (Mérai et al., 2014), which causes loss of centromere identity and dispersion of the H3K9me2 marked centromeric heterochromatin (Schoft et al., 2009). Moreover, expression of the DNA glycosylase DEMETER (DME) in the VN causes demethylation of some genes and transposable elements (TEs) in the CG sequence context (Ibarra et al., 2012; Park et al., 2017; Schoft et al., 2011). This active demethylation process is required to induce the expression of a small number of genes involved in pollen-tube function, which are critical for male fertility (Borg et al., 2021a; Khouider et al., 2021). On the other hand, the biological function of the VN chromatin decompaction is still debated, and two possible models have been proposed. The first hypothesis is that global decondensation occurs to deliberately reactivate TEs, to allow the production of small RNAs that are then transferred from the VC to the SCs to reinforce silencing in the germline (Ibarra et al., 2012; Martínez et al., 2016; Slotkin et al., 2009). Alternatively, chromatin decondensation could allow the reactivation of the thousands of copies of ribosomal RNA genes that are normally heterochromatinized and segregated in the chromocenters (Mérai et al., 2014). Indeed, the VC is extremely metabolically active, as it needs to rapidly elongate the pollen tube to find the ovule, which makes it one of the fastest growing eukaryotic cells known (Johnson et al., 2019).

We recently discovered that two Arabidopsis methyl-reader proteins, Methyl-CpG-binding domain 5 and 6 (MBD5 and MBD6, or MBD5/6 for short), redundantly silence both genes with promoter methylation and TEs (Ichino et al., 2021). MBD5/6 bind CG methylated DNA and represses transcription by recruiting the J-domain protein SILENZIO (SLN). While most genes and TEs repressed by DNA methylation are bound by MBD5 and MBD6, only a subset of them are derepressed in *mbd5/6* mutants (Ichino et al., 2021). This led us to investigate what makes these loci sensitive to the loss of the methyl-readers and whether the derepression might be limited to certain cell types. Here we show that loss of silencing in *mbd5/6* and in *sln* occurs specifically in the VN of developing pollen grains. Given the special epigenetic state of this cell, we propose that the molecular function of MBD5/6 is revealed during pollen development because of diminished chromatin compaction, which makes the VN particularly prone to loss of silencing. Consistent with this, we found that the *mbd5/6* phenotype is enhanced in leaves by mutation of the linker histone H1, which causes chromatin decompaction in leaves (Choi et al., 2020; Rutowicz et al., 2019). This result highlights the importance of evolving several redundant layers of silencing mechanisms to ensure the maintenance of TE repression. Furthermore, this work provides a comprehensive single-nucleus RNA-seq dataset of Arabidopsis developing pollen nuclei, which has allowed novel insights into the transcriptome plasticity of male gametophytes.

## RESULTS

### The *mbd5/6* TE derepression phenotype is strongest in developing pollen

We previously observed that in inflorescence tissue the *mbd5/6* and *sln* mutant plants show a mild derepression of a small number of TEs and genes that are regulated by DNA methylation, among which a clear example was the gene *FWA* (Ichino et al., 2021). *FWA* is normally methylated and silent in all tissues except for the endosperm, a tissue that surrounds and nourishes the embryo, where only the maternal copy is demethylated, thus allowing its monoallelic expression (Kinoshita et al., 2004). A very small loss of methylation in the *FWA* promoter is also observed in the pollen VN (Schoft et al., 2011). Consistently, recently published RNA-seq datasets detected low levels of *FWA* expression in microspores and bicellular pollen (Julca et al., 2021) (Figure S1). Given the specificity of the *FWA* expression pattern in reproductive tissues and given that our previously published *mbd5/6* and *sln* RNA sequencing (RNA-seq) experiments were performed with unopened flower buds, we wondered whether the loss of silencing occurred only in specific cell types within the flower. To investigate this, we performed manual dissections of unopened flower buds to separate the anthers, which contain the developing pollen grains, from the rest of the flower bud, including carpels, petals, and sepals (named “no-anthers” for short) (Figures 1A and S1B). We then performed RNA-seq to compare the gene expression profiles in wild-type, *mbd5/6* and *sln* (Table S1). We found that *FWA* derepression in *mbd5/6* and in *sln* occurred exclusively in the anthers (Figure 1B). Furthermore, for both genotypes, differential gene expression analysis detected, in anthers, more than 200 upregulated and less than 40 downregulated transcripts (Figure 1C), which is consistent with the previously described function of this methyl-reader complex in gene silencing (Ichino et al., 2021). Of note, only 34 upregulated transcripts were detected in the no-anthers fraction of *mbd5/6* (10 in *sln*) (Figure 1C). Therefore, while we cannot rule out that MBD5/6 could regulate transcription in some rare cell types within the no-anthers fraction, the majority of the derepression signal measured in flower buds derives from the anther tissue.

**Figure 1:**
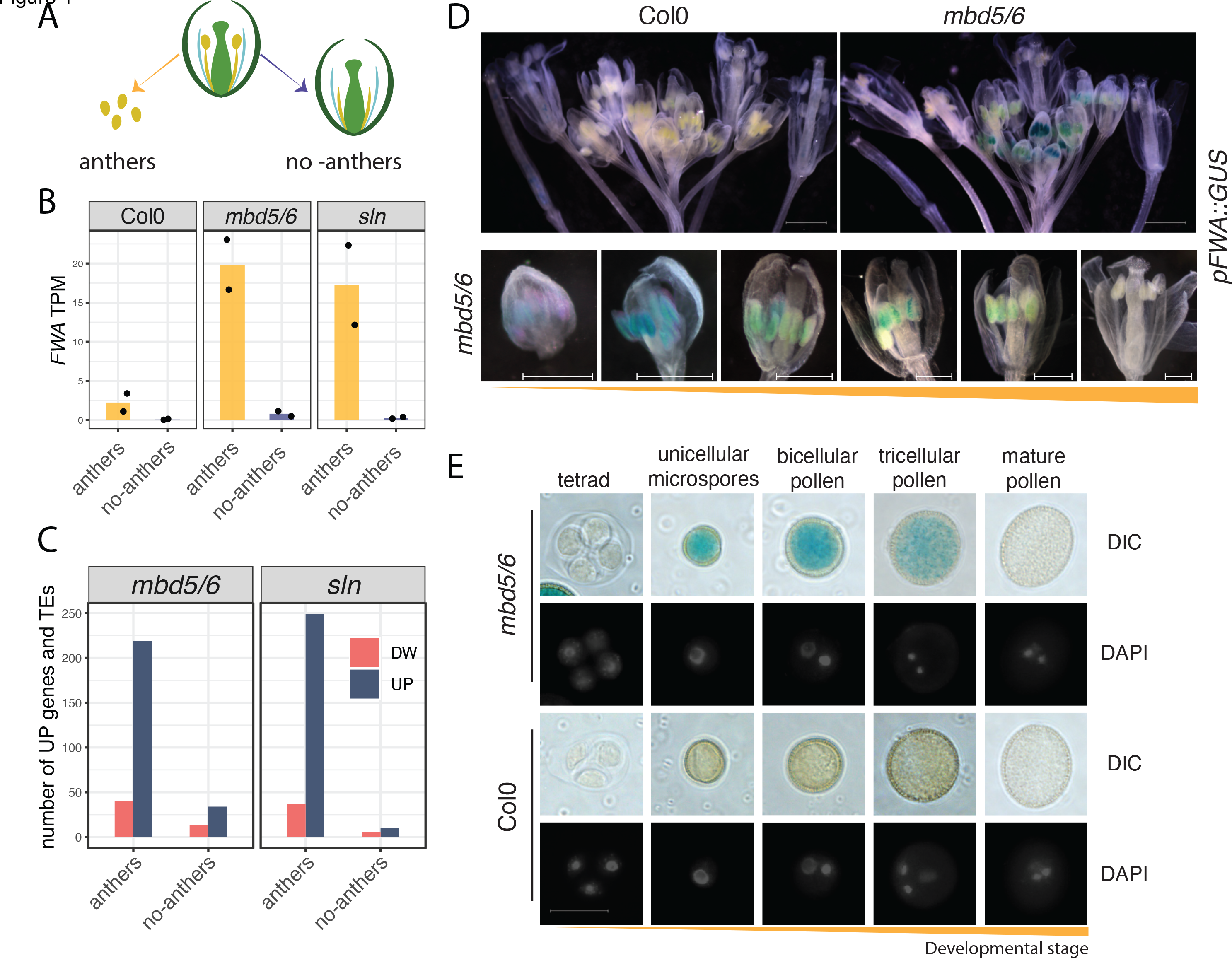
*FWA* derepression in *mbd5/6* is restricted to developing pollen grains. A) Cartoon depicting the dissection strategy: anthers were dissected from unopened flower buds and separated from the rest of the sample (“no-anthers”), which includes sepals, petals, and carpels. B,C) RNA-seq analysis of the dissected fractions. Panel B shows *FWA* expression levels (transcripts per million [TPM]), panel C shows the number of up- or downregulated transcripts for each mutant relative to wild-type controls. D) Microscopy images of GUS stained and cleared pFWA::GUS lines in the Col0 (wild-type) or *mbd5/6* backgrounds (representative of more than 10 lines per genotype). Scale bars upper panels: 1000 μm; lower panels: 500 μm. E) Images of male gametophytes at the indicated developmental stages, isolated from *pFWA::GUS* lines in the Col0 (wild-type) or *mbd5/6* backgrounds. Scale bars: 20 μm.

To confirm this result with an orthogonal approach, we generated transcriptional reporter lines in which the *FWA* promoter drives the expression of the *beta-GLUCURONIDASE* (*GUS*) gene (Ikeda et al., 2007) in the wild-type or *mbd5/6* genetic backgrounds. As previously showed, in wild-type plants we detected GUS staining only in young developing seeds, which contain the endosperm tissue, where *FWA* is endogenously expressed (Figure S1C). Instead, in *mbd5/6* mutant plants we detected a clear speckled pattern in the anthers at different developmental stages (Figure 1D). The intensity of the GUS precipitate appeared strongest in early developmental stages and then tended to fade away in the most mature anthers (Figure 1D). Further inspection confirmed that the GUS staining corresponded to the developing male gametophytes within the anther locules (Figure 1E). To understand which stages of pollen development display *FWA* expression, we performed DAPI staining of developing spores from the reporter lines. We found that GUS positive pollen grains corresponded to all stages of development, from the uninuclear microspores to the bicellular and tricellular pollen grains, but the signal was not detectable anymore in mature tricellular pollen (Figure 1E). Overall, these data indicate that within the anther tissue, *FWA* is reactivated only in the male gametophytes, from early stages of pollen development, and its expression decreases in mature pollen. Given that pollen mitotic divisions occur over a timeframe of about three days (Alvarez-Buylla et al., 2010; Sanders et al., 1999), it is possible that *FWA* mRNA expression decreases before the loss of GUS protein, as we further address below.

*FWA* derepression only in the male gametophyte was unexpected because MBD5, MBD6 and SLN are all broadly expressed across many different tissues (Figure S1). Therefore, their expression pattern cannot explain the pollen specificity of their phenotype. Moreover, in *Arabidopsis thaliana*, DNA methylation typically represses a largely overlapping set of genes and TEs in different tissues. The *FWA* gene, for instance, is very highly expressed both in vegetative and in reproductive tissues in *met1-3* mutant plants, in which CG methylation is lost genome-wide because of a mutation in the maintenance *DNA METHYLTRANSFERASE 1* (*MET1*) gene (Figure S2A).

The specificity of *FWA* derepression in *mbd5/6* for developing pollen grains prompted us to investigate and compare the gene expression patterns in seedlings, unopened flower buds, and mature pollen (see Methods and Figure S2B and S2C for the mature pollen collection procedure). We detected 17 upregulated DEGs in *mbd5/6* seedlings, 57 in flower buds, and 176 in mature pollen (Figures 2A, S2D). Only 3 out of 17 seedlings up-DEGs were upregulated in flower or pollen, while 33 out of 57 flower up-DEGs were upregulated in pollen, but only one of them in seedlings (Figures S2E and S2F). Similar results were obtained with *sln* (Figures S2D and S2G). When we plotted the distribution of the expression fold-changes in the different tissues, we noticed that the pollen DEGs tend to have a positive fold-change in flower buds as well, although lower in magnitude compared to the fold-change in pollen (Figures 2B and S2H). On the other hand, these loci are not upregulated in seedlings (Figures 2B and S2H). This is different from what is observed in the methylation mutant *met1*, in which about half of the pollen DEGs are upregulated in seedlings as well (Figures 2B). Therefore, a portion of the *mbd5/6* pollen DEGs are repressed by DNA methylation in seedlings, but loss of MBD5/6 alone is not sufficient to reactivate them in that tissue.

**Figure 2:**
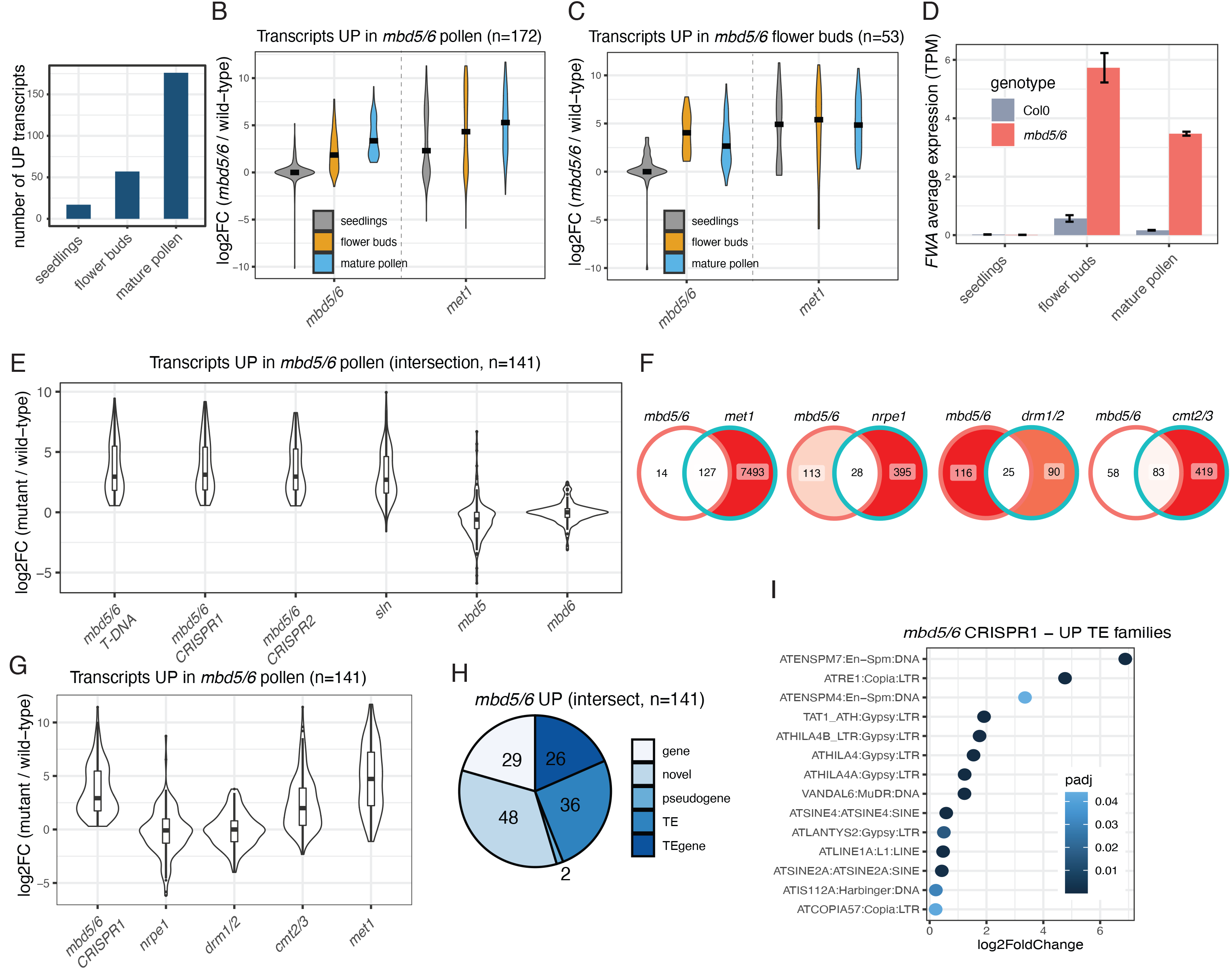
Tissue specificity of the *mbd5/6* transcriptional derepression phenotype. A) RNA-seq analysis of *mbd5/6* versus wild-type, performed with the indicated tissues. B,C) Distribution of the RNA-seq log2 fold-change (log2FC) for the transcripts (genes and TEs) that are upregulated in *mbd5/6* mature pollen (B) or unopened flower buds (C). Black line indicates the median. The *met1* seedlings RNA-seq dataset is from (Stroud et al., 2012) and the *met1* flower buds RNA-seq is from (Ichino et al., 2021). D) Expression of *FWA* in Col0 and *mbd5/6*, from the RNA-seq datasets shown in A. E,G) Comparison of the mature pollen RNA-seq pattern in different mutants. The overlayed violin plots and boxplots show the distribution of the log2 fold-changes of the upregulated transcripts that overlap between the three different *mbd5/6* alleles. F) Overlaps between upregulated DEGs in mature pollen in the indicated mutants. H) Features of the *mbd5/6* mature pollen upregulated transcripts. The “novel” transcripts were identified via a transcript reannotation based on all the mature pollen RNA-seq datasets included in this study (see Methods and Figure S3). I) Analysis of significantly upregulated TE families in *mbd5/6* (mature pollen).

We also noticed that a small number of genes, including *FWA,* are more strongly upregulated in *mbd5/6* flower buds than in mature pollen (Figures 2C, 2D, S2D, and S2I). This is consistent with the observations made with the *pFWA::GUS* reporter line, which showed loss of GUS signal in mature pollen (Figures 1C and 1D).

### The transcripts upregulated in *mbd5/6* pollen are genes and TEs repressed by CG methylation

Given that the mature pollen RNA-seq allowed detection of a higher number of DEGs compared to the previously published dataset (Ichino et al., 2021), we sought to investigate the features of these transcripts in greater depth. While inspecting the genome browser tracks, we noticed that several loci with aligned reads did not correspond to any annotated gene or transposable element (Figure S3). In order to be able to detect these loci as DEGs, we performed a reannotation of transcripts based on our Col0, *mbd5/6*, *sln*, and *met1* mature pollen RNA-seq datasets (see Methods and Figure S3). This allowed us to not only detect more DEGs but also refine the existing annotations to obtain more accurate transcriptional start, end, and splicing sites (Figure S3). We then used these annotations to analyze a mature pollen RNA-seq dataset that includes the *mbd5* and *mbd6* single mutants, three different *mbd5/6* double mutants, *sln*, *met1,* the RdDM mutants *drm1 drm2 (drm1/2)* and *nrpe1*, which lose non-CG methylation at RdDM sites, and the *cmt2 cmt3 (cmt2/3)* mutant, which affects methylation at heterochromatic TEs (Stroud et al., 2014).

We obtained about 200 upregulated transcripts in the *mbd5/6* samples and in *sln*, while less than 20 were detected in the *mbd5* and *mbd6* single mutants (Figures 2E, S4A and S4B, Table S2). This is consistent with our previously published observation that MBD5 and MBD6 are genetically redundant, and that SLN is the repressor acting downstream of the methyl-readers (Ichino et al., 2021). The *mbd5/6* up-DEGs (n=141, intersection between the three *mbd5/6* mutants) were mostly a small subset of the *met1* up-DEGs (127/141) and overlapped well with the *cmt2/3* up-DEGs (83/141), but not as much with the DEGs of the RdDM mutants *nrpe1* and *drm1/2* (28/141, 25/141) (Figure 2F). Consistently, these genes had an overall positive fold-change in *met1* and *cmt2/3*, but not in *nrpe1* and *drm1/2* (Figures 2G). This indicates that the MBD5/6 targets are mostly heterochromatic loci, as previously observed in flower buds (Ichino et al., 2021), and while some of them, like *FWA*, are euchromatic and methylated by the RdDM machinery, they are not upregulated in RdDM mutants, but only in *met1*, suggesting that they are mainly repressed by CG methylation.

The *met1* up-DEGs included both loci that are promoter methylated and unmethylated, and the latter are likely indirect targets (Figure S4C). In contrast, almost all the *mbd5/6* upregulated DEGs were promoter methylated and were not expressed in wild-type flowers (Figures S4C and S4D). Consistently, we found that a vast majority of them were TEs, TE-genes (transposon related genes as defined in the TAIR10 genomic annotations), or novel transcripts obtained with our pollen reannotation pipeline (Figure 2H). TE family analysis showed that the MBD5/6 targets include both retrotransposons and DNA transposons, both of which are upregulated in *met1* as well (Figures 2I, S4E and S4F) (Quesneville, 2020). Therefore, MBD5 and MBD6 do not regulate a specific class of TEs, but broadly regulate TE families that are repressed by CG methylation.

We also identified a few functional genes that, similarly to *FWA*, have promoter methylation and are repressed by MBD5/6 and MET1 (Figure S4G). One of them was ANTAGONIST OF LIKE HETEROCHROMATIN PROTEIN 2 (ALP2), a gene domesticated from a transposon, which antagonizes the function of POLYCOMB REPRESSIVE COMPLEX 2 (PRC2) (Velanis et al., 2020).

Overall, these datasets show that among the investigated tissues, the transcriptional phenotype of *mbd5/6* and *sln* is strongest in pollen, and that the MBD5/6 targets are genes and TEs repressed by CG methylation. Given that pollen grains contain different nucleus types, we next sought to pinpoint in which specific nuclei the transcriptional changes occurred during pollen development, using a single-nucleus resolution approach.

### Single nucleus RNA-seq reveals the transcriptional landscape of developing male gametophyte nuclei

Transcriptomic analysis of Arabidopsis pollen has been previously performed either by isolation of discrete developmental stages of whole pollen grains with a Percoll step-gradient (Honys and Twell, 2004; Julca et al., 2021) or via Fluorescence-Activated Nuclei Sorting (FANS) of VN and SN from mature pollen (Borg et al., 2021a). It has been shown that the expression profile of whole mature pollen samples strongly correlates with that of the mature VN, and not with the SN, because most of the pollen mRNA content derives from the cytoplasm of the VC (Borg et al., 2021a). Therefore, analysis of staged pollen fractions does not allow direct investigation of the GN or SN transcriptomes, and doesn’t provide a complete and continuous picture of the transcriptional changes that occur during the development of microspore nuclei (MN) and of the VN lineage. For these reasons we developed a method to capture the transcriptome of each nucleus of the developing pollen using the 10X genomics platform (Figure 3A). Briefly, we isolated male gametophytes from Arabidopsis inflorescences by gentle homogenization of the tissue, and we purified the mixed spores from contaminating tissue by centrifugation over a Percoll cushion. Visual inspection of the sample confirmed that this method allowed the isolation of mixed-stage pollen grains, from the uninuclear microspores to the tricellular mature pollen (Figure 3B). To break the pollen wall and release the nuclei, we vortexed the sample intermittently in the presence of glass beads, as previously done for VN and SN isolation (Borges et al., 2012; Santos et al., 2017). We then sorted the nuclei based on DAPI to purify them from the debris (Figure S5A, S5B, and S5C), and we performed single-nucleus RNA-seq with the 10X Genomics Chromium platform. The pre-processing and quality control (QC) of the datasets were done with Seurat using standard workflows (methods section “Analysis of snRNA-seq”, Table S3).

**Figure 3:**
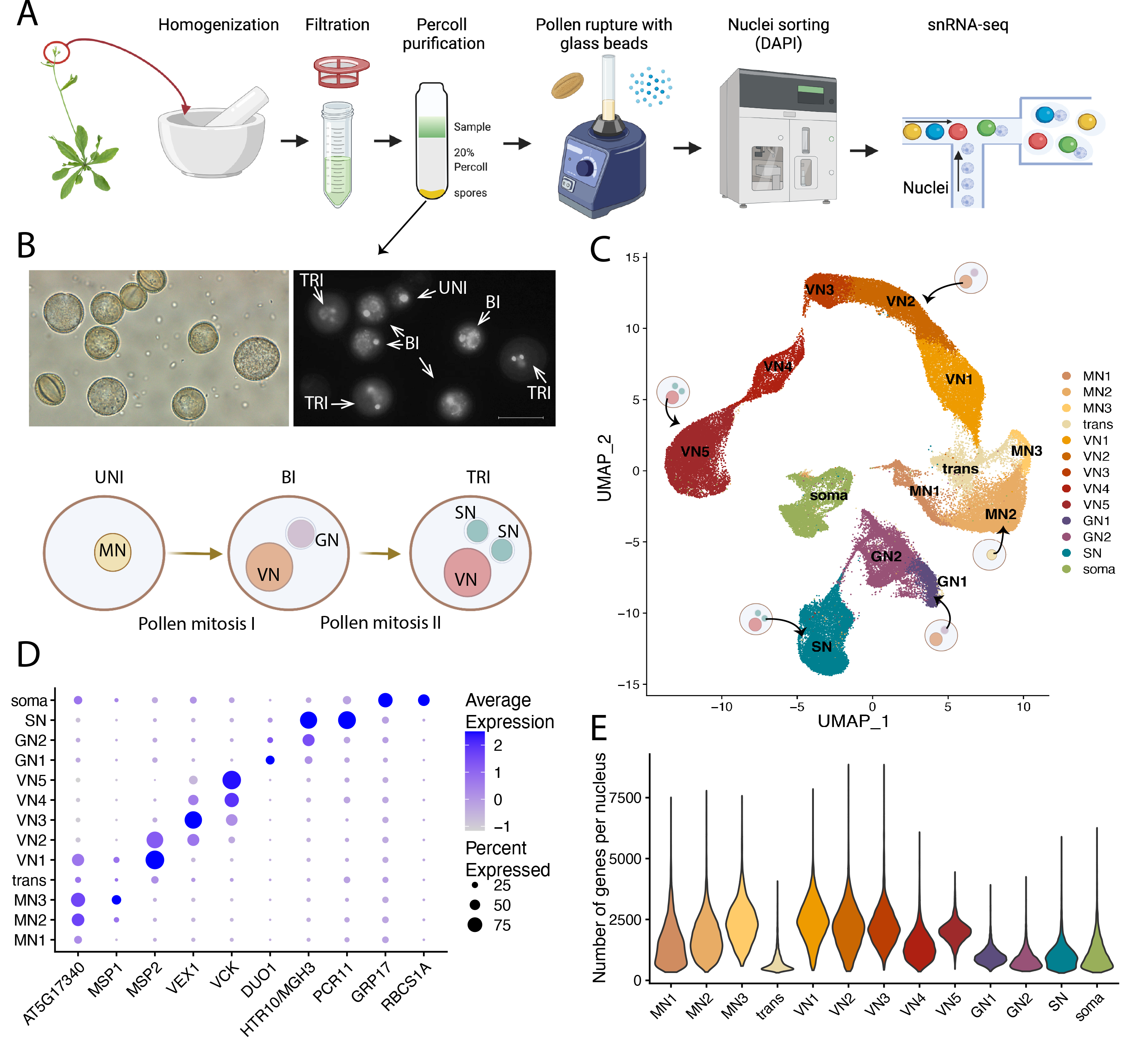
Single-nucleus RNA-seq of developing male gametophytes. A) Cartoon representation of the protocol developed for this study. B) Upper panels: representative image of the mixed spores sample obtained after gradient centrifugation. Left image: DIC, right image: DAPI. Scale bar: 20 μm. Lower panel: cartoon representation of pollen development. “Uni”, “bi”, and “tri” indicate the uninuclear, bicellular, and tricellular pollen stages. C) UMAP of all integrated snRNA-seq datasets with clusters annotations. D) Dot plot showing the cluster specificity of the expression of known markers. Dot size: percentage of cells in which the gene was detected. Dot color: scaled average expression. E) Violin plot showing the distribution of the number of unique genes detected per nucleus, in each cluster.

To visualize the data, we performed dimensionality reduction via Uniform Manifold Approximation and Projection (UMAP) followed by unsupervised clustering (Figure 3C). Inspection of several well-known markers of the different pollen nucleus types allowed the identification of clusters corresponding to MN, VN, GN, SN, and some contaminating nuclei (labeled “Soma”) likely deriving from somatic tissues that surround the developing pollen grains in the anthers (Figures 3C and 3D). We identified marker genes for each cluster, and the complete list is available in Table S4. The cluster identity assignments were also confirmed by selecting the top 20 markers for each cluster, and plotting their normalized expression in several RNA-seq datasets from the EVOREPRO database which were generated from bulk RNA-seq of whole pollen grains at different developmental stages (Julca et al., 2021) (Figure S5D).

To put the expression defects of *mbd5/6* and *sln* in the context of development, we first sought to characterize the wild-type expression patterns of all nucleus types in this dataset. The next section of the manuscript describes the nuclei clusters, before delving into the transcriptional changes found in *mbd5/6* and *sln*.

We observed that the VN nuclei were distributed in the UMAP along a clear developmental trajectory, proceeding from the bicellular pollen stage to the tricellular and mature pollen (labeled as VN1 to VN5) (Figure 3C). This was indicated by the initial expression of *MICROSPORE-SPECIFIC PROMOTER 2* (*MSP2*) which is a VN marker of the early bicellular stage (Honys et al., 2006), followed by the tricellular stage VN marker *VEGETATIVE CELL EXPRESSED 1* (*VEX1*) (Engel et al., 2005), and then by the expression of *VEGETATIVE CELL KINASE 1* (*VCK1*), which is strongest in the latest stages of mature pollen (Grant-Downton et al., 2013) (Figure 3D). Based on this observation, we generated a developmental trajectory using Monocle3 (Cao et al., 2019) to rank the VN nuclei according to their predicted pseudotime, which represents the amount of progress that each nucleus has made towards differentiation (Figures S5E and S5F). We then identified all the genes that change as a function of pseudotime (see Methods), and performed hierarchical clustering to find patterns, which revealed different waves of transcriptional regulation (Figure S5G and Table S5). After performing gene ontology (GO) analysis we found that that the genes that were expressed in the early stages and then gradually downregulated were enriched in several GO terms related to mRNA processing and ribosome biogenesis, the genes that were transiently upregulated in the middle of the trajectory were related to Golgi vesicle transport and endocytosis, and the genes upregulated in the later stages were associated with cell tip growth and pollen tube development GO terms (Figures S5G and Table S5). This analysis suggests that VN nuclei initially work towards building the machinery required for extensive transcription and translation, and later use it to produce the proteins needed for membrane expansion and pollen tube growth, which is an extremely demanding metabolic process entailing extensive membrane trafficking (Johnson et al., 2019). The high transcriptional activity of these nuclei is revealed also by the higher average number of genes detected compared to the SN and GN clusters (Figure 3E).

We then identified the GN nuclei through the master transcription factor *DUO POLLEN 1* (*DUO1*), which is expressed in the GN right after pollen mitosis I (Borg et al., 2009, 2011) (Figure 3D). The SN nuclei instead were marked by sperm cell specification genes that are activated by DUO1, such as *MGH3/HTR10* and *PCR11* (Borg et al., 2011). While *PCR11* is known to be sperm cell specific, *MGH3/HTR10* was shown to be expressed in the GN of the late bicellular pollen as well as in the SN (Okada et al., 2005), and indeed we were able to detect strong expression of this gene in the GN2 cluster (Figure 3D). Several other known DUO1 targets were also detected in GN2 and SN clusters (Figure S5H) (Borg et al., 2011). The relative expression patterns of these genes suggest that the GN2 cluster comes after GN1 in the developmental timeline (Figures S5H and S5I). Gene ontology analysis of the GN cluster markers highlighted terms related to cell cycle and chromosome organization (Table S6).

We identified 3 clusters corresponding to microspore nuclei (MN1, MN2, and MN3). These clusters express the previously described microspore specific marker *AT5G17340* (da Costa-Nunes, 2013), while *MICROSPORE SPECIFIC PROTEIN 1* (*MSP1*) (Honys et al., 2006) was only detectable in MN2 and MN3 (Figure 3D). The expression pattern of these two genes suggests that these nuclei are distributed along a developmental trajectory that proceeds from MN1 to MN3. Consistently, an analysis of known cell cycle markers revealed that the MN1 cluster is enriched in S-phase markers, while the MN3 cluster is enriched in M-phase markers, which supports the directionality of the trajectory (Figure S5J). Indeed, MET1, which is highly expressed in S-phase to methylate the newly synthesized DNA (Du et al., 2012), was also highly expressed in MN1 (Figure S5K). Gene ontology analysis revealed that the MN3 cluster, in addition to cell cycle terms, was also enriched in terms related to ribosome biogenesis, suggesting that late microspores are already beginning to establish a transcriptional program aimed at increasing the metabolic output of the upcoming VN (Table S6).

The UMAP analysis identified a group of cells localizing between the MN and the VN1 clusters, that despite having a gene expression profile similar to the one of MN cells, were characterized by very low numbers of genes detected per nucleus and very few cluster markers (Figures 3C, 3E, Table S3, and Table S4). We named this cluster “transitory” (“trans”) because we envision that it could correspond to cells that are in the process of undergoing mitotic division, and therefore might be going through a transient global decrease in transcriptional output (Palozola et al., 2017). However, we acknowledge that these cells could also correspond to damaged nuclei or chromatin fragments, and therefore we did not include this cluster in our downstream analyses.

We noticed that one of the top markers for the MN clusters was *DEMETER-LIKE PROTEIN 3* (*DML3*) which is a poorly characterized DNA demethylase enzyme (Table S4 and Figure S5K). This prompted us to investigate the family of Arabidopsis demethylases. As expected, we observed a strong enrichment of *DEMETER* (*DME*) in the early VN (Figure S5K), which is consistent with DME’s known role in demethylating some genes that are required for pollen tube growth (Khouider et al., 2021; Park et al., 2017). ROS1 (DML1) has also been shown to participate in the process of demethylation in the VN, and consistently we observed that its expression was strongest in late MN and VN1. DML3 instead was strongly expressed in the MN1 and MN2 clusters, and interestingly DML2 was mostly enriched in the GN1 cluster (Figure S5K). This suggests that different demethylases might have cell type-specific functions during pollen development.

We next inspected the expression patterns of the different DNA methyltransferases and we observed that CMT1 was only expressed in GN1, while its homologue CMT3, which has CHG methyltransferase activity, was expressed in MN and SN as well as GN (Figure S5K). CMT2 instead, which has CHH specificity, was most highly expressed in early VN, and therefore it is likely responsible for the known increase in CHH methylation levels in the VN, given that the other CHH methyltransferases, DRM1 and DRM2, were very lowly expressed (Figure S5K) (Calarco et al., 2012; Ibarra et al., 2012). Lastly, MET1, as previously mentioned, has an expression pattern that strongly correlates with cell cycle, being most highly expressed in MN1, GN, and SN.

Inspection of the machinery required for the deposition and removal of histone marks revealed that some enzymes were broadly expressed while others were restricted to specific lineages (Figure S5K). Of note, the H3K27 methyltransferases *CURLY LEAF* (*CLF*) and *SWINGER* (*SWN*) were enriched in the VN and depleted from SN, which is consistent with a recent report of the erasure of H3K27 in sperm nuclei (Borg et al., 2020). We also looked at the expression patterns of the small RNA machinery, given its prominent role in pollen biology (Borges et al., 2011; Oliver et al., 2022). As previously described, we found that AGO5 was expressed in the GN and AGO1 in microspores and VN (Figure S5K).

The role of histone variants in shaping the epigenome of male gametophytes has gained increasing attention in recent years (Borg et al., 2021b). In addition to *MGH3/HTR10,* which is a well-known sperm specific H3 variant (H3.10), we validated the sperm specificity of *H2B.8* (*HTB8*), which was recently found to mediate chromatin condensation in sperm to reduce nuclear size (Buttress et al., 2021; Jiang et al., 2020) (Figure S5L). Interestingly, *H2B.5* and *H2B.7* were enriched in sperm as well, and these variants haven’t been functionally analyzed yet (Figure S5L). *H2B.10* was instead preferentially expressed in the late VN, as previously reported (Jiang et al., 2020). It will be interesting to determine whether H2B.10 contributes to chromatin relaxation in the VN.

Overall, our single nucleus RNA-seq dataset provides a comprehensive overview of gene expression profiles throughout the development of the Arabidopsis male gametophytes and constitutes a source for future studies. We next used this dataset to investigate at single nucleus resolution the transcriptional changes occurring in *mbd5/6* and *sln*.

### The MBD5/6 targets are derepressed in the VN lineage

To investigate the specificity of the *mbd5/6* derepression during pollen development, we performed the snRNA-seq experiment with Col0, *mbd5/6*, and *sln* mutant plants (Figure 4A). We first looked at the *FWA* gene and we found that its expression in *mbd5/6* begins increasing in the late microspore stage, peaks in the VN of bicellular pollen, and then starts decreasing in the late bicellular stage of the VN development (Figure 4B). This is consistent with the results obtained with the *pFWA::GUS* reporter line, and it suggests that GUS signal in early tricellular pollen may be in part due to persistent protein expression when the mRNA levels are decreased (Figures 1C and 1D). *FWA* expression was very mild in *mbd5/6* GN and SN, suggesting that its upregulation is mostly restricted to the MN/VN lineage.

**Figure 4:**
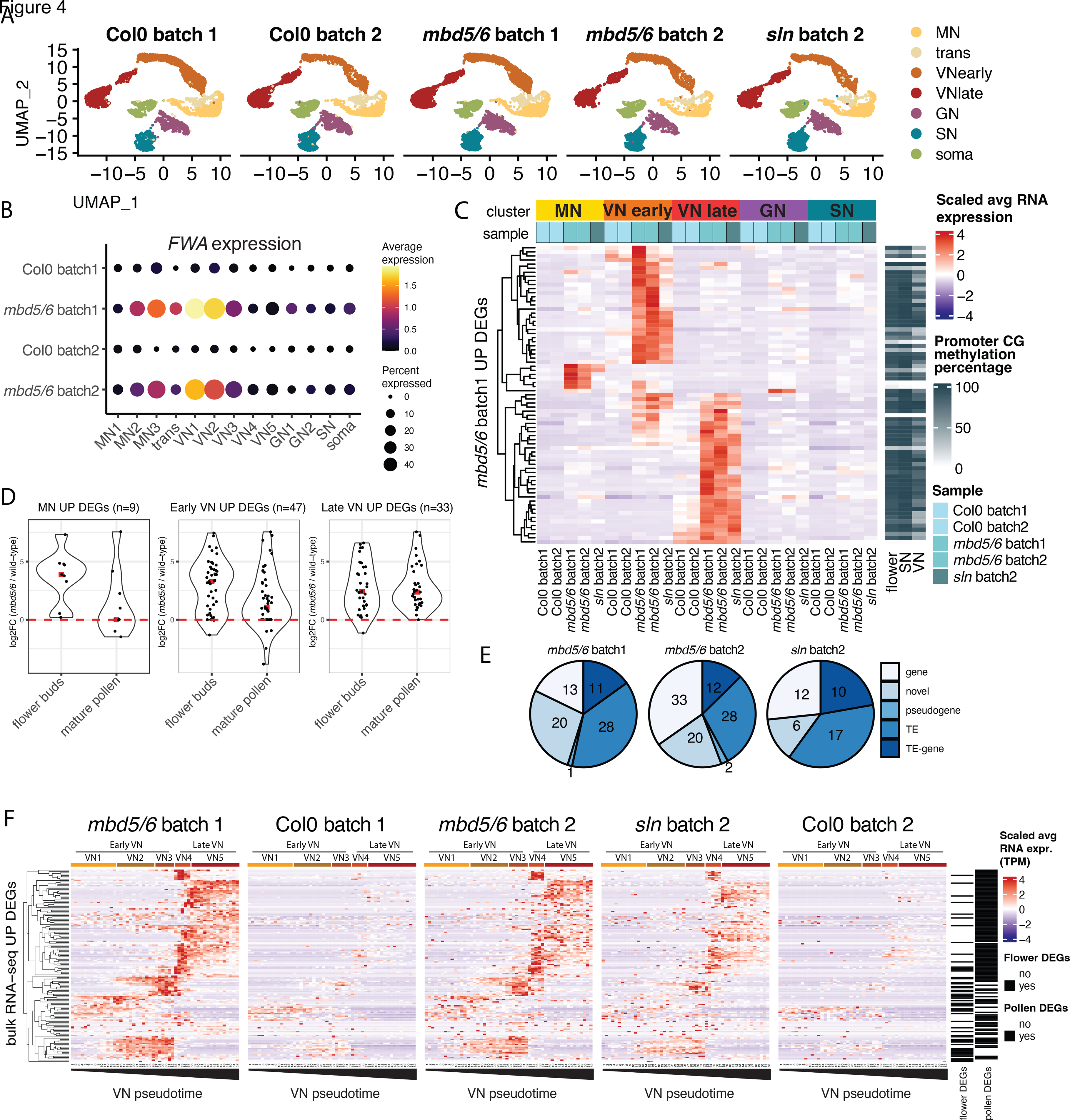
Transcriptional derepression in *mbd5/6* is limited to the MN/VN lineage. A) UMAP plots of the integrated Col0, *mbd5/6*, and *sln* snRNA-seq datasets. “Batch 1” and “Batch 2” indicate two independent experiments. B) Dot plot of *FWA* expression in Col0 and *mbd5/6*, at each cluster. Dot size: percentage of cells in which *FWA* was detected. Dot color: scaled average expression. C) Heatmap representation of the *mbd5/6* batch 1 upregulated transcripts (union of all cluster). Shown is the snRNA-seq scaled average expression level per cluster, in the indicated samples. Right columns: wild-type CG methylation percentage at promoters. The VN and SN BS-seq data is from (Ibarra et al., 2012). D) Violin plot showing the expression change in *mbd5/6* vs Col0 (log2 fold-change) obtained by bulk RNA-seq in flowers or mature pollen, for the indicated groups of genes (red dash: median). The MN DEGs are the union of MN1, MN2, and MN3 DEGs, the early VN DEGs are the union of VN1, VN2, and VN3, the late VN DEGs are the union of VN4 and VN5 DEGs. “DEGs” always indicates all transcripts (genes and TEs). E) Classification of the upregulated transcripts for each experiment (union of all clusters). F) Heatmap of scaled snRNA-seq expression along the VN pseudotime trajectory (see Figure S5E). Shown is the union of flower bud and mature pollen *mbd5/6* upregulated transcripts obtained by bulk RNA-seq (see Figure S2).

To test whether other MBD5/6 targets showed the same pattern as *FWA*, we performed a differential gene expression analysis comparing *mbd5/6* to wild-type for each cluster, including samples from two independent experiments. We employed a stringent approach for calling DEGs to limit the false positives, as explained in the Methods and in Figure S6. We observed that the clusters corresponding to the same nucleus type (such as VN4 and VN5) had a similar pattern of differential expression (Figure S6F), therefore we decided to group together these clusters for ease of visualization, and only display the MN, early VN, late VN, GN, and SN groups in a scaled heatmap (Figures 4A and 4C). We observed that a small number of transcripts are upregulated in microspores, and most of the *mbd5/6* and *sln* differentially expressed genes (DEGs) are specific either to the early or late VN clusters, while they are not strongly upregulated in GN or SN (Figures 4C and S6). To validate this observation, we analyzed the expression of these DEGs in the bulk RNA-seq datasets performed on unopened flower bud tissue and on mature pollen. We reasoned that the microspore and early VN DEGs should be more strongly upregulated in the “unopened flower bud” bulk RNA-seq sample, which contains anthers with developing pollen grains, compared to the mature pollen sample. Indeed, the result confirmed the prediction and validated the stage specificity of the DEGs (Figure 4D). As expected, these loci were characterized by promoter methylation and included several TEs and novel annotations (Figures 4C and 4E). The number of DEGs obtained from the snRNA-seq data was lower than that obtained from the bulk RNA-seq, likely because of the high variability in single cell data (see Methods and Figure S6). However, our approach was able to identify high confidence and reproducible DEGs, including *FWA* and several TEs.

We next sought to investigate the dynamics of expression of the DEGs along the VN developmental trajectory. We plotted the expression levels of the loci that were defined as upregulated in *mbd5/6* by mature pollen or flower buds bulk RNA-seq (n=171) (Figure 4F). Although most of these loci were not called as snRNA-seq DEGs based on our stringent snRNA-seq analysis pipeline (129/171, Figure S6G), 137 out of 171 were detected in the single-nucleus data and showed upregulation in *mbd5/6* or *sln* compared to the controls (Figure 4F). We observed a progressive upregulation along the trajectory, with 54 genes being expressed in early VN and then silenced, and 83 genes being upregulated in the late VN (Figure 4F). Interestingly, this same pattern, but with much lower magnitude, was observed in the wild-type controls as well, whereby genes upregulated early in *mbd5/6* or *sln* also tended to be detected in the early stages in WT, and the late upregulated genes in the later stages. This suggests that the stage specificity of the derepression likely reflects the expression of needed developmental stage specific transcription factors.

Overall, the snRNA-seq data revealed that loss of silencing in *mbd5/6* and *sln* begins in the late microspore stage and then becomes progressively more prominent in the VN along its developmental trajectory, while GN and SN are not affected.

### Other silencing mutants show broad derepression across all pollen nucleus types

Intrigued by the VN specificity of the MBD5/6 targets, we wondered whether mutations in other genes involved in silencing of methylated DNA might show a similar or different pattern. To address this, we performed male gametophyte snRNA-seq with a panel of other well characterized mutants that have TE derepression. We selected two “strong” mutants characterized by an extensive loss of methylation and high numbers of derepressed TEs, *met1* and *ddm1*, and two “weak” mutants that have fewer derepressed TEs, and very limited loss of methylation, *mom1* and *morc1,2,4,5,6,7 hextuple* (hereafter named “*morc*”). As previously noted, *met1* displays a genome-wide loss of CG methylation and limited loss of non-CG methylation, with strong TE derepression, while mutations in the nucleosome remodeler *DECREASE IN DNA METHYLATION 1* (*DDM1*) cause loss of CG and non-CG methylation only in pericentromeric heterochromatin, and strong derepression of heterochromatic TEs (Jeddeloh et al., 1999; Lippman et al., 2004). Mutations in *MORPHEUS’ MOLECULE1* (*MOM1*) instead cause a mild derepression of heterochromatic TEs with very limited loss of DNA methylation, but the exact mechanism of MOM1-mediated silencing is not understood (Amedeo et al., 2000; Han et al., 2016). Similarly, mutating the entire family of MORC GHKL ATPases (*morc1,2,4,5,6,7* hextuple mutant) causes derepression of some heterochromatic TEs that do not display a strong loss of methylation (Harris et al., 2016). We investigated each mutant with a matched wild-type control (Figure 5A). In the *met1* sample we obtained relatively fewer cells corresponding to the most mature stages of pollen development (VN4, VN5, SN) probably due to the phenotypic defects in flower development in the *met1* plants (Figure 5A and Table S2).

**Figure 5:**
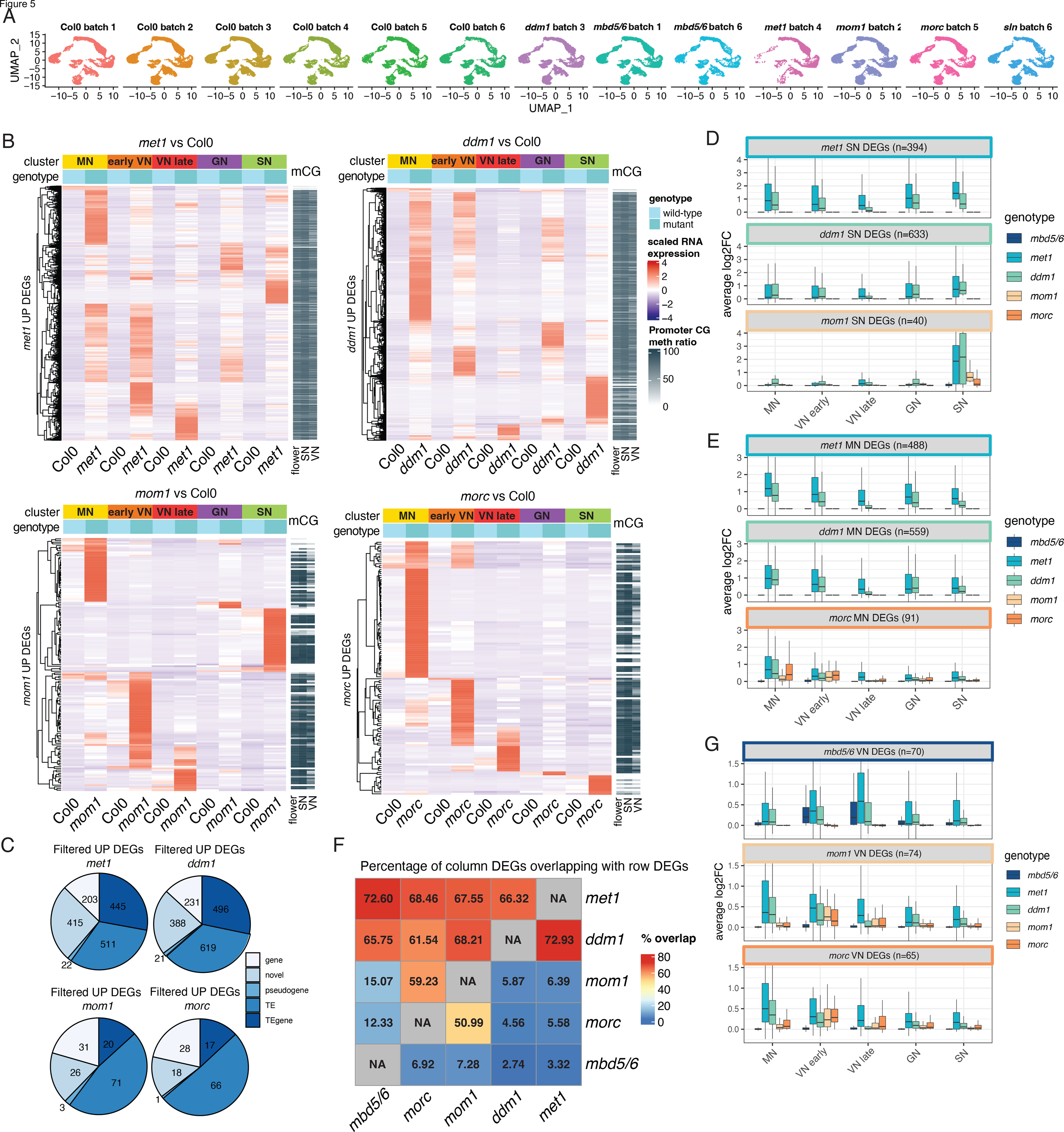
*met1*, *ddm1*, *mom1*, and *morc* mutants display loss of silencing in all pollen nucleus types. A) UMAP plots of all the snRNA-seq datasets used in this study. B) Heatmaps of the genes and TEs upregulated in the indicated mutants (union of all clusters). Shown is the snRNA-seq scaled expression level of the cluster averages in the indicated samples. Right columns: wild-type CG methylation percentage at the promoters. The VN and SN BS-seq data is from (Ibarra et al., 2012). C) Classification of the upregulated transcripts for each mutant (union of all clusters). D,E,G) Boxplots of the average log2 fold-change difference (mutant over its matched wild-type control) for the indicated groups of genes. F) Heatmap representation of the percentage of DEGs upregulated in the mutant indicated in the column label (“column DEGs”) that overlap with DEGs upregulated in the mutant annotated in the row label (“row DEGs”). For instance, 72.6% of the *mbd5/6* upregulated DEGs overlap with the *met1* upregulated DEGs. “DEGs” always indicates all transcripts (genes and TEs).

In all cases, we detected DEGs distributed among all clusters (Figures 5B and S7). For all mutants, about two thirds of the DEGs were TEs or novel transcripts, characterized by promoter methylation (Figures 5B and 5C). The heatmap visualization of the expression of these DEG among the different cluster groups highlighted how, unlike *mbd5/6* and *sln*, these mutants showed transcripts strongly upregulated in every cell type in developing and mature pollen (Figures 5B and S7). The *morc* mutant was particularly enriched in microspore-specific DEGs, suggesting that MORC proteins could play an important role in the early stages of pollen development. The *mom1* mutant instead was relatively more enriched in SN specific DEGs compared to the other mutants. We noticed that while the *ddm1* and *met1* SN DEGs were similarly upregulated in all clusters, the *mom1* SN DEGs were instead more strongly upregulated in SN compared to the other clusters, even in *ddm1* and *met1* (Figure 5D). This suggests that in weaker mutants like *mom1*, only the loci that have strong, cluster specific promoters tend to be derepressed. Consistently, the *morc* MN DEGs also had a clear cluster specificity, because even in *ddm1* and *met1* they were more strongly upregulated in MN compared to the other clusters (Figure 5E).

We next compared the lists of upregulated DEGs of the different mutants, and we observed that while a large proportion of the MBD5/6 targets were upregulated in *met1* and *ddm1*, they were not upregulated in *mom1* nor in *morc* (Figure 5F). Similarly, the MOM1 and MORC targets were mostly a subset of the MET1 and DDM1 targets, and they largely overlapped with each other, but were not upregulated in *mbd5/6* (Figure 5F). Consistently, when plotting the distribution of the fold-changes we found that the *mbd5/6* VN DEGs were not upregulated in *mom1* nor in *morc,* while the *mom1* and *morc* VN DEGs were not upregulated in *mbd5/6* (Figure 5G). Therefore, the MBD5/6 targets constitute a unique subset of the loci regulated by DNA methylation. This prompted us to further investigate the features of these loci with the goal of understanding why this is the only mutant that displays a strong VN specificity.

### Loss of histone H1 uncovers a derepression phenotype of *mbd5/6* and *sln* **in non-reproductive tissues.**

The VN is a large nucleus and adopts a very peculiar chromatin state, characterized by global decondensation and loss of heterochromatic chromocenters (He et al., 2019; Mérai et al., 2014). We wondered whether this unusual relaxed chromatin state could be related to the VN specificity of the loss of silencing in *mbd5/6*. During germline development, the process of decondensation begins in the late microspore stage, when the linker histone H1 is depleted, likely to facilitate the relaxation of chromatin (He et al., 2019). The loss of H1 has also been proposed to facilitate access of the demethylase DME to the promoters of some genes and TEs, which is one of the reasons why CG methylation levels in the VN are diminished (Ibarra et al., 2012). We reasoned that this unique chromatin state might make the VN particularly sensitive to loss of silencing in *mbd5/6*. Of note, the *mbd5/6* derepression also began in the late microspore stage, thus correlating with the timing of the H1 depletion from this nucleus (He et al., 2019).

Given that only a small subset of the transcripts that are derepressed in *met1* are reactivated in *mbd5/6* as well, we sought to test whether these loci could be characterized by higher levels of accessibility. We took advantage of a recently published dataset of Assay for Transposase-Accessible Chromatin (ATAC-seq) performed on nuclei isolated from wild-type GN, SN, and mature VN (Borg et al., 2021a). We observed that in the wild-type VN there was a clear increase in chromatin accessibility at the MBD5/6 targets but not at the MET1-specific targets (which are not upregulated in *mbd5/6*) (Figure 6A). This observation supports the hypothesis that the MBD5/6 targets might be particularly prone to loss of silencing in the VN because they gain accessibility in this cell type. The MBD5/6 targets also tended to lose more CG methylation in the VN, compared to the MET1 targets, possibly because the openness of their promoters facilitates access of DME, as previously suggested (Figure 6B) (He et al., 2019). However, while the DME-mediated demethylation of specific pollen fertility genes in the VN is very extensive, causing these genes to be highly expressed and not regulated by MBD5/6, the loss of methylation at the MBD5/6 targets was milder, and only led to very low levels of expression in the wild-type VN (Figures 6B-6E).

**Figure 6:**
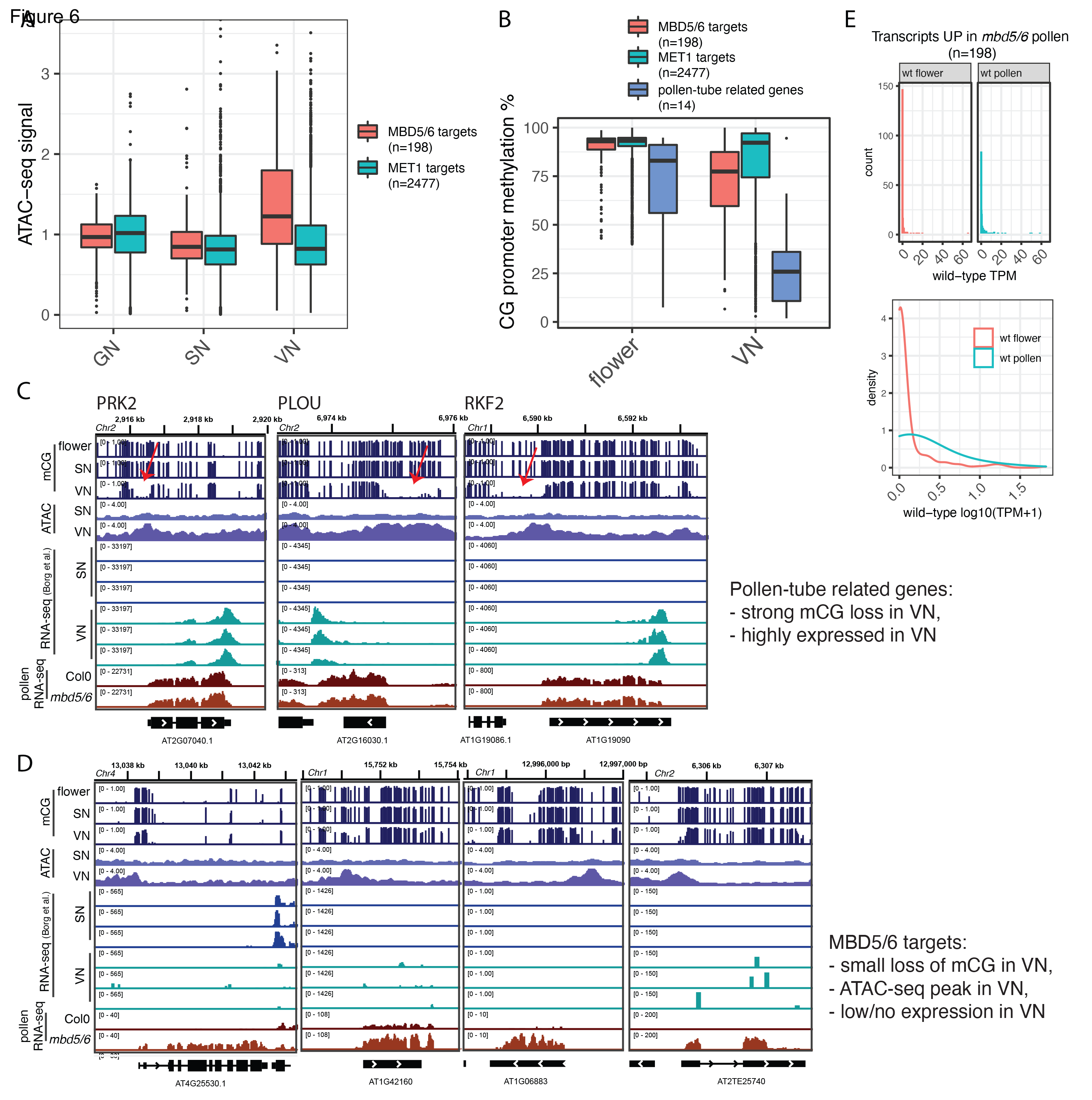
The MBD5/6 targets are characterized by increased accessibility in the wild-type VN. A) Boxplot showing the average ATAC-seq signal at the promoters of the MBD5/6 targets (loci with promoter methylation upregulated in *mbd5/6* mature pollen RNA-seq) or MET1 targets (loci with promoter methylation, upregulated in *met1* but not in *mbd5/6* mature pollen). The VN, GN, and SN ATAC-seq data is publicly available (Borg et al., 2021a). B) Boxplots of average CG promoter methylation percentage at the loci defined in A and at 14 manually curated pollen-tube related genes that are demethylated by DME in the VN (list in Methods) (Borg et al., 2021a; Khouider et al., 2021). The SN and VN BS-seq data is from (Ibarra et al., 2012). C-D) Genome browser tracks showing examples of genes required for pollen fertility (C) (Khouider et al., 2021), and genes/TEs repressed by MBD5/6 (D). The SN and VN RNA-seq and ATAC-seq data is publicly available (Borg et al., 2021a). The scale for ATAC-seq is kept constant in all screenshots for ease of comparison. The red arrows point to the strong loss of CG methylation that allows expression of these genes in the wild-type VN. E) Histograms of the wild-type expression level (TPM) in unopened flower buds and mature pollen, for the loci derepressed in *mbd5/6*. The lower plot shows the smoothened trend of the data to highlight a mild increase in expression of these transcripts in pollen.

This analysis suggested that the reason why there isn’t any detectable TE derepression in *mbd5/6* and *sln* seedlings could be that chromatin compaction compensates for the loss of the methyl-readers and is sufficient to maintain gene silencing. To test this idea, we crossed *mbd5/6* and *sln* with the *h1.1/h1.2* double mutant (hereby named *h1* for short), which has been shown to have decondensed chromatin in seedlings (Choi et al., 2020; He et al., 2019; Rutowicz et al., 2019). After performing RNA-seq in seedlings we found that the *FWA* gene, which is not expressed in *mbd5/6* or *sln* seedlings, was very mildly expressed in *h1*, and clearly enhanced in *mbd5/6 h1* and in *sln h1* (Figure 7A). At a global level, we also observed an enhancement of derepression of methylated loci in the higher order mutants compared to *h1, mbd5/6,* or *sln* alone, indicating that when chromatin compaction is impaired as in *h1*, the function of MBD5/6 and SLN is revealed in seedlings (Figures 7B-7D). We found a group of about 50 genes that were significantly upregulated only in *mbd5/6 h1* or *sln h1* (Figure 7C). Interestingly, these sites also show positive, though milder, upregulation in *mbd5/6* and *sln* alone (Figure 7E). Therefore, they tend to be mildly upregulated in *mbd5/6* even in the presence of H1, but their expression is increased when H1 is depleted. These results suggest that MBD5/6 and SLN have gene silencing functions in a broader range of tissues, but that redundancy with other silencing pathways prevents upregulation of target genes in most cells other than the VN.

**Figure 7:**
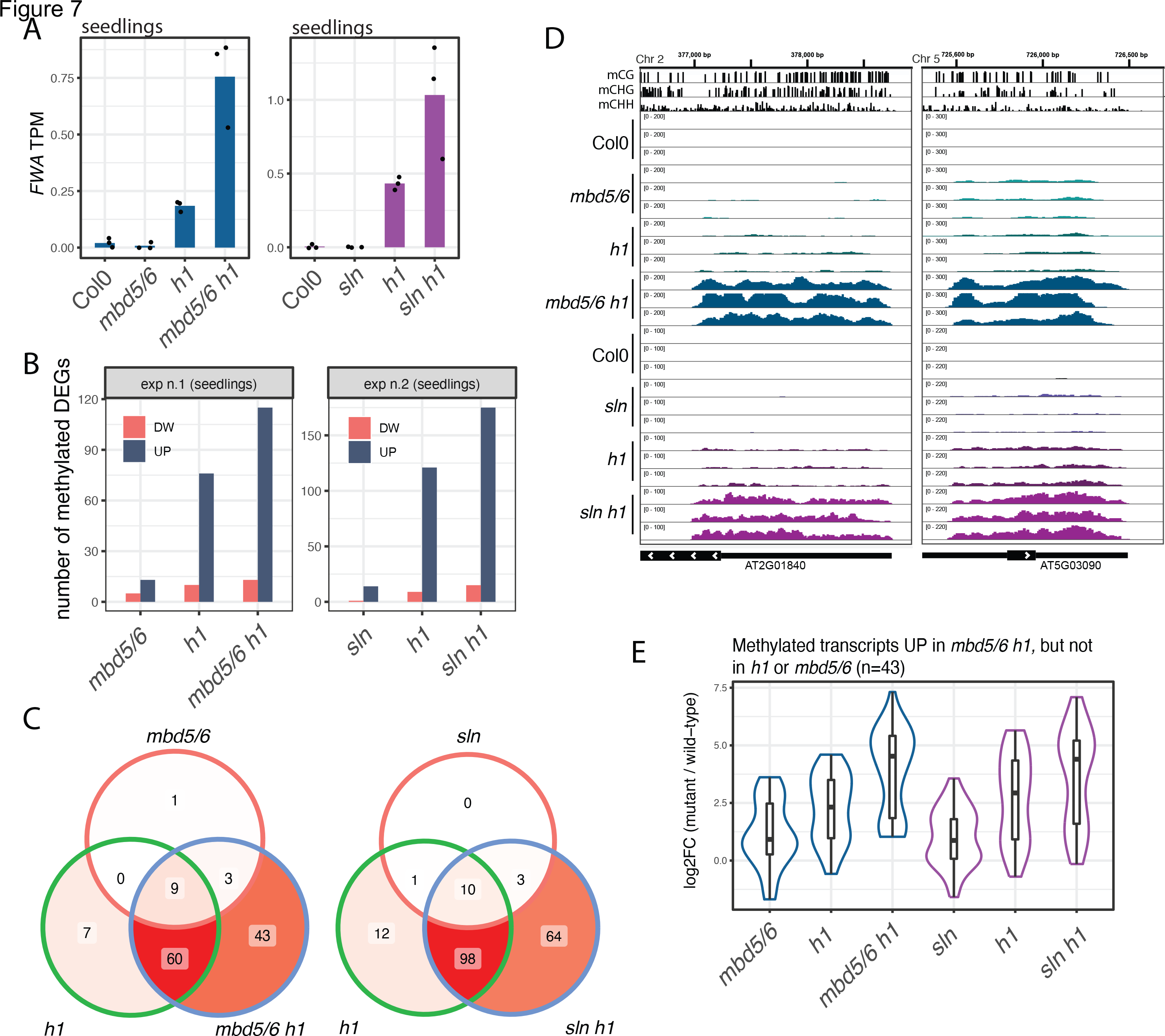
H1 mutation enhances the *mbd5/6* derepression phenotype in seedlings. A) Barplots of average *FWA* expression in seedlings in the indicated genotypes (seedlings RNA-seq). The dots correspond to individual replicates. B) Number of up- or downregulated transcripts for each mutant relative to wild-type controls. Only the loci with average CG promoter methylation higher than 40% are shown. C) Venn diagrams of the upregulated methylated transcripts. D) Examples of two loci that are enhanced by combined loss of MBD5/6 and H1 (BS-seq: flower buds, RNA-seq: seedlings). E) Overlay of violin plots and boxplots showing the distribution of log2 fold-changes for the indicated transcript group.

## DISCUSSION

The complexity of gene silencing mechanisms reflects a strong evolutionary pressure to preserve genome integrity by repressing mobile elements in eukaryotic organisms. The redundancies between different silencing pathways make it challenging to study these mechanisms; indeed, a role of Arabidopsis methyl-reader proteins in gene silencing downstream of DNA methylation has only been discovered recently (Ichino et al., 2021). In the present manuscript we show that the function of MBD5, MBD6 and SILENZIO becomes evident in the vegetative nucleus of pollen, a specific cell type that has diminished function of important silencing factors that normally maintain chromatin compaction. To facilitate an extremely high transcriptional and metabolic activity, the vegetative cell undergoes dramatic chromatin decondensation, which makes it vulnerable to increased transposon activity. Our results suggest that the MBD5/6 methyl-readers are important to combat the upregulation of transposons that become accessible to the transcriptional machinery in this fragile, but reproductively crucial, cell type.

In addition to TEs, a limited number of functional genes were found to be upregulated in *mbd5/6,* and these genes are not known to be involved in pollen development (Figure S4G). Indeed, we did not observe any pollen developmental defect in *mbd5/6*. Strong TE derepression instead could interfere with the biological activities of the VN by, for instance, sequestering energy from other important cellular processes required for pollen tube development. Future studies aimed at investigating pollen function and fertilization could reveal functional defects in *mbd5/6* pollen. The *met1* mutant instead had a much stronger transcriptional deregulation phenotype and displayed a decrease in the relative number of mature pollen grains, which could be due to strong transposon derepression or to the deregulation of specific genes. It is likely that combined mutations of multiple members of the MBD protein family could lead to a stronger phenotype, more closely resembling that of *met1*.

Our results do not exclude the possibility that MBD5 and MBD6 could play important roles in other rare cell types. For instance, the central cell of the female gametophyte shares several features with the pollen VN: decondensed and partially demethylated chromatin due, in part, to DME’s demethylation activity (Park et al., 2016; Pillot et al., 2010). While thousands of pollen grains are present in each flower bud, only about 50 central cells exist at the right developmental stage. Therefore, the investigation of transcriptional changes occurring in central cells requires direct isolation of this rare cell type. The endosperm tissue, which originates from fertilization of the central cell, has decondensed chromatin as well, and it is the tissue in which imprinting occurs in Arabidopsis (Gehring and Satyaki, 2017). The *FWA* gene for instance becomes demethylated in the central cell, and later on, in the endosperm, is expressed only from the demethylated allele inherited from the central cell (Kinoshita et al., 2004). Given the role of MBD5/6 in silencing *FWA* in pollen, these repressors could maintain silencing of the paternal and methylated allele of *FWA* in endosperm, to ensure imprinted expression. It will be interesting therefore to investigate whether they play a broad role in imprinting regulation.

We found that other repressors that act downstream of DNA methylation, such as MOM1 and the MORC proteins, do not show the same VN-specific phenotype as MBD5/6. Consistent with this finding, mutations in these factors cause reactivation of TEs in non-reproductive tissues such as seedlings (Amedeo et al., 2000; Harris et al., 2016). The observation that in *mom1* and *morc*, which are mild mutants like *mbd5/6*, TE derepression is not enhanced in the pollen VN suggests that the tissue specificity of the *mbd5/6* transcriptional phenotype could be due to the specific mechanism of silencing of the methyl-readers as opposed to the other repressors. MBD5/6 could be required for gene silencing only at accessible promoters by, for instance, preventing the transcriptional machinery from binding the DNA. Indeed, the MBD5/6 targets gain ATAC-seq peaks in the wild-type VN but remain very lowly or not expressed (Figure 6). On the other hand, if the silencing mechanism of MOM1 and MORC involves chromatin decompaction, as previously suggested for MORC proteins (Guillaume et al., 2012), this might explain why the MBD5/6 targets, which are already decompacted, are not derepressed in *mom1* and *morc* (Figure 5G). Future experiments are needed to investigate these hypotheses regarding the molecular mechanisms of action for these silencing factors.

Most of the *morc* DEG were strongly derepressed in microspore nuclei. This is consistent with the observation that several members of the MORC family are most strongly expressed in these nuclei, while MOM1 for instance has a broad expression pattern and has DEGs in all nuclei types (Figure S5M). MORC5 is highly expressed in GN1 (Figure S5M), but we observed only one upregulated DEG in that cluster (Figure S7). It is possible that MORC5 plays a role in the GN that is not related to gene or TE repression. The strong transcriptional effect of MORC proteins in early male gametogenesis is reminiscent of the known role of some animal MORC family members in the male germline (Kojima-Kita et al., 2021; Pastor et al., 2014; Shi et al., 2018; Weiser et al., 2017). However, plant MORC proteins are not required for male fertility, and we did not observe any pollen development defect in *morc hextuple* mutant, thus leaving open the question of whether Arabidopsis MORCs play a role in pollen biology.

In addition to providing novel insights into silencing mechanisms, this work contributes a comprehensive snRNA-seq dataset capturing the developing male gametophyte nuclei throughout development, from early microspores to mature pollen. This transcriptional atlas allowed us to make several novel discoveries that suggest new avenues of research, and constitutes an important resource that can be further explored to gain insights into pollen biology and development. For instance, some epigenetic factors like MORC5, DML2, and CMT1, which were poorly characterized due to their low expression levels in most tissue, were found to be highly expressed in GN1, a cell type for which a transcriptome hasn’t been generated yet. This dataset can be easily explored with an interactive website (https://singlecell.mcdb.ucla.edu/snRNAseq_pollen/), and the pollen transcriptome reannotations and codes used to generate them are available for download on Github (https://github.com/clp90/mbd56_pollen/).

## LIMITATIONS OF THE STUDY

While our study identifies an important role for MBD5 and MBD6 in the pollen VN, we cannot exclude the possibility that these proteins could play important functions in other rare cell types that we haven’t directly inspected so far, such as the central cell or the endosperm.

## Supporting information

Table S7 10X DEGs - Related to Figure 4 and 5

Table S6 GO all clusters - Related to Figure 3

Table S5 GO VN lineage - Related to Figure 4

Table S4 cluster markers - Related to Figure 3

Table S3 cellranger and DoubletFinder - Related to Figure 3

Table S2 bulk pollen DEGs - Related to Figure 2

Table S1 all genomics datasets-Related to Figures 1,2,6,7

## ACKNOWLEDGMENTS

We thank Dr. Lachezar Nikolov and Nathan Tran for their help with the GUS staining, Drs. Yoko Ikeda and Takashi Araki for the pFWA::GUS plasmid, Michael Borg and David Twell for advice on pollen protocols, Giuseppe Barisano and Elena Brivio for advice on 10X sample preparation and data analysis, Brandon Boone, Jake Harris and members of the Jacobsen lab for insightful discussion, the UCLA Technology Center for Genomics & Bioinformatics (TCGB) core for the 10X libraries, the Eli and Edythe Broad Center of Regenerative Medicine and Stem Cell Research University of California, Los Angeles Flow Cytometry Core Resource for sorting, the UCLA Broad Stem Cell Research Center BioSequencing core for high throughput sequencing. This work was supported by a Philip Whitcome Pre-Doctoral Fellowship in Molecular Biology to L.I., a Ruth L. Kirschstein National Research Service Award (F32GM136115) to C.L.P., a George G. & Betsy H. Laties Graduate Fellowship in Molecular Plant Biology to S. W., and NIH R35 GM130272 to S.E.J.. S. E. J. is an Investigator of the Howard Hughes Medical Institute. Graphical illustrations in Figure 3 were created with BioRender.com.

## AUTHOR CONTRIBUTIONS

L. I. and S. E. J. conceived the study and designed the research; L.I. performed the experiments and analyzed the data; C. L. P. and M. C. contributed to the data analysis;

J. Y., Y. X. and S. W. contributed to experiments; R. K. P. contributed to the experimental design and discussions; L.I. drafted the manuscript; S. E. J. and C. L. P. contributed to manuscript writing; all authors discussed and reviewed the manuscript.

## STAR METHODS

### RESOURCE AVAILABILITY

#### Lead contact

Further information and requests for resources and reagents should be directed to and will be fulfilled by the lead contact, Steve Jacobsen.

#### Materials availability

Seeds for all Arabidopsis lines generated in this study are available upon request to the lead contact.

#### Data and code availability

All sequencing data have been deposited at GEO and are publicly available as of the date of publication. Accession numbers are listed in the key resources table

(GSE202422, reviewers access token: ilkdeawifrcnxkh). The snRNA-seq data can be freely inspected in an interactive website (https://singlecell.mcdb.ucla.edu/snRNAseq_pollen/).

Microscopy data reported in this paper will be shared by the lead contact upon request.

This paper analyzes existing, publicly available data. These accession numbers for the datasets are listed in the key resources table.

The pollen transcriptome reannotations and the code used to generate them have been deposited in github at https://github.com/clp90/mbd56_pollen. DOIs are listed in the key resources table. All other codes used in this study are available upon request.

Any additional information required to reanalyze the data reported in this paper is available from the lead contact upon request.

### EXPERIMENTAL MODEL AND SUBJECT DETAILS

All plants used in this study were in the Columbia-0 ecotype (Col-0) and were grown on soil in a greenhouse under long-day conditions (16h light / 8h dark). The experiments performed on seedlings were done after growing the plants on 1/2 MS medium plates under constant light.

The following mutant lines were obtained from Arabidopsis Biological Resource Center (ABRC) or previously generated as indicated in the references: *mbd5*

(SAILseq_750_A09.1), *mbd6* (SALK_043927), *sln* (SALK_090484) (Ichino et al., 2021), *met1-3* (CS16394), *nrpe1-11*, *drm1 drm2* (Chan et al., 2006), *cmt2 cmt3* (Stroud et al., 2014), *ddm1-2* (Vongs et al., 1993), *mom1-3* (SALK_141293), *morc* hextuple consisting of *morc1-2* (SAIL_893_B06) *morc2-1* (SALK_072774C) *morc4-1* (GK-249F08) *morc5-1* (SALK_049050C) *morc6-3* (GABI_599B06) and *morc7-1* (SALK_051729) (Harris et al., 2016). The *h1.1-1 h1.2-1* double mutant (referred to as *h1*) consists of SALK_128430 and GABI_406H11 (Zemach et al., 2013). The *mbd5/6* T-DNA double mutant and the *mbd5/6* CRISPR-1 mutant were previously described (Ichino et al., 2021). The *mbd5/6* CRISPR- 2 mutant was generated via CRISPR/Cas9 in the Col0 background as described in the next section.

The *mbd5/6 h1* and *sln h1* mutants were generated by crossing. The seedlings used for RNA-seq of *sln*, *h1*, *sln h1* and wild-type controls where F3 plants derived from individual F2 segregants of the cross with the indicated genotypes. The *mbd5/6 h1* experiment instead was performed with Col0, *mbd5/6,* and *h1* controls grown from the original batch of seeds used for the cross.

### METHOD DETAILS

#### Generation of *mbd5/6* CRISPR-2 line

The *mbd5/6* CRISPR-2 line was generated using the previously published pYAO::hSpCas9 system (Yan et al., 2015). We cloned two different guides for MBD5 (G1: ACCGGAGAACCCGGCTACTC, G2: GAAATCTAAAGTTCGATGTG) and two different guides for MBD6 (G1: TTCCGGTGCCACAGCTGGTT, G2: ATATGTTAGGGTTACTTAAT) performing four sequential cloning steps. Each guide was cloned in the AtU6-26-sgRNA cassette by overlapping PCR with primer tails containing the guide sequence. The PCR products were cloned into the SpeI site of the pYAO::hSpCas9 destination plasmid in four steps of SpeI digestion followed by In-Fusion (Takara). The final vector was electroporated into AGLO agrobacteria and transformed in Col0 plants by agrobacterium-mediated floral dipping. The transgenic lines were genotyped both by PCR amplification of the area surrounding the two guides to detect large deletions, and by Sanger sequencing to detect indels. The *mbd5/6* CRISPR-2 line was isolated in the T3 generation and was confirmed to have segregated out the pYAO::hSpCas9 transgene. This line has a homozygous G insertion at the MBD6 guide 1 region, which causes a frameshift and a premature STOP codon. The MBD5 mutation instead is a large inversion encompassing the entire region between the guide 1 and guide 2, which causes a frameshift and a premature STOP codon.

#### Generation of transgenic lines

The *pFWA::GUS* transgenic lines were generated with a published expression vector (Ikeda et al., 2007). The vector was electroporated into AGL0 agrobacteria that were used for plant transformation by agrobacterium-mediated floral dipping. Transformants were selected 1/2 MS medium plates with kanamycin, and the kanamycin resistant T1 plants were transplanted on soil. The GUS staining and imaging was performed in the T1 generation.

#### GUS staining and imaging

The experiment was performed in multiple batches with a total of least 10 individual *pFWA::GUS* T1 transgenic lines for each genotype (Col0 and *mbd5/6* T-DNA). The two genotypes were always processed side by side. One inflorescence from each plant was clipped and placed in cold acetone on ice. The samples were transferred to -20°C for 30 min. After the incubation, the inflorescences were washed two times with room temperature water, and then transferred in GUS staining solution (1 ml of 0.1 M X-Gluc, 1.71 ml of 1 M Na2H, 0.79 ml of 1 M NaH2, 2.5 ml of 0.1 M potassium ferrocyanide, 2.5 ml of 0.1 M potassium ferricyanide, 100 ul of Triton X100, water up to 50 ml). The samples were vacuum infiltrated for about 10 minutes and then incubated at 37°C for about 4-5 hours. The reaction was stopped by transferring the samples to 70% ethanol, and the inflorescences were kept in 70% ethanol for 3 days with gentle shaking, changing the ethanol solution every day. Next, the samples were washed with water and then incubated in ClearSee solution for one day (Kurihara et al., 2015). Samples were imaged with a ZEISS Stemi 508 Stereo Microscope with Axiocam 208 Color. The images were processed with ZEISS ZEN lite.

For high magnification imaging of GUS-stained developing pollen grains, five to seven inflorescences were harvested in 500-700 μl of 0.1 M mannitol solution, in 1.5 ml tubes. The flower buds were gently disrupted with a pestle to release the spores in solution. The solution was filtered over 80 μm nylon mesh two times, and then centrifuged over a cushion of 500 μl of 45% Percoll in 0.1 M mannitol, for 10 min at 800 g. The pellet containing the cleaned mixed stage spores was resuspended in 200 μl of GUS staining solution and incubated at 37°C for about 4-5 hours. Next, the samples were collected by brief centrifugation at 500 g, and the pellets were resuspended in 10 μl of DAPI buffer (0.4 μg/ml DAPI solution in 0.1 M sodium phosphate buffer, 10 mM EDTA-disodium salt and 0.1% Triton X-100, pH 7.0) (Park et al., 1998). After a 5 min incubation, 5 μl of each sample were transferred to a microscopy slide, covered with a glass coverslip and sealed with nail polish. Samples were imaged immediately with an Axio Imager.D2 upright microscope.

#### Dissection of anthers for RNA-seq

Anthers were manually dissected from 20 individual flower buds (stage 10-12) for each sample (two samples per genotype obtained from different plants). Dissected anthers were placed immediately in 350 μl of ice-cold RLT buffer from the RNeasy micro kit (Qiagen, 74004) keeping the tube on ice. For each flower bud, all other remaining tissues were placed in a separate tube with 350 μl of ice-cold RLT buffer, to prepare the “no-anthers” fraction. The tissues were disrupted on ice in the RLT buffer using a sterile pestle (Axygen, PES-15-B-SI). We then proceeded with RNA extraction using the RNeasy micro kit (Qiagen, 74004) starting from step 4 (addition of one volume of 70% ethanol). In-column DNAse digestion was carried on following manufacturer’s instructions. RNA- seq libraries were generated using the TruSeq Stranded mRNA Library Prep Kit (Illumina) with the standard workflow, starting with 300-400 ng of RNA.

#### Bulk RNA-seq of seedlings, flower buds, and mature pollen

Samples:

Biological triplicates were generated for each genotype.

Seedlings RNA-seq was done by harvesting ∼10-15 14 days old seedlings from ½ MS plates and flash freezing them in liquid nitrogen (replicated were grown in separate plates).

Unopened flower bud RNA-seq was done by harvesting one inflorescence (excluding open flowers) from an individual plant for each sample and freezing it immediately in liquid nitrogen.

Mature pollen RNA-seq was performed by harvesting the pollen with the previously described vacuum method (Johnson-Brousseau and McCormick, 2004). Briefly, about 150 plants were grown for each genotype. We assembled an in-house filtering system on a vacuum cleaner to place three different nylon meshes at the end of the tube in this order: 80 μm, 31 μm, 7 μm. Using this system, we aspirated the pollen from the plants. The 80 μm and 31 μm filters block the flower parts such as petals, the soil and other particulate, while the pollen accumulates on the 7 μm mesh. We then collected the pollen from the 7 μm nylon mesh using a pipette with 0.1 M mannitol, and transferred it into a 1.5 ml tube. The solution was centrifugated for 5 minutes at 500 g. The supernatant was removed, and the pollen pellet was frozen in liquid nitrogen and stored at -80°C. This procedure was repeated every 2/3 days to obtain replicates for each genotype.

In the case of *met1*, given the difficulty to obtain high numbers of homozygous mutant plants, we used a different protocol to purify mature pollen from a smaller number of plants. We harvested ∼500 μl of open flowers from *met1* and Col0 control plants in 2- ml protein low bind tubes (Eppendorf). We then added 800 μl of Galbraith buffer (45 mM MgCl2, 30 mM C6H5Na3O7.2H2O [Trisodium citrate dihydrate], 20 mM MOPS, 0.1% [v/v] Triton X-100, pH 7), and vortexed the samples for 3 minutes at max speed to release the pollen from the anthers. The suspension was filtered with an 80 μm nylon mesh into a new 1.5 ml tube. The procedure was repeated one more time with the same flowers to increase the yield of pollen. The two aliquots of filtered pollen suspension were combined and centrifuged for 5 min at 800 g, 4°C. The pollen pellet was frozen in liquid nitrogen and stored at -80°C. This procedure was repeated every 2/3 days to obtain replicates.

RNA extraction and library preparation:

Frozen samples were disrupted with a tissue grinder and RNA extraction was performed with the Zymo Direct-zol RNA MiniPrep kit (Zymo Research), or in the case of the pollen samples with the QIAGEN RNeasy Mini kit (Qiagen). In both cases, the in-column DNase digestion was performed.

RNA-seq libraries were generated using the TruSeq Stranded mRNA Library Prep Kit (Illumina), following the manufacturer’s instructions and starting with 1 µg of RNA as input for flowers and seedlings, and 300-500 ng for the pollen samples.

#### Single-nucleus RNA-seq of developing male gametophytes

The protocol for isolation of mixed-stage male gametophytes was developed starting from a published protocol (Dupl’Akova et al., 2016). We processed 2 or 3 genotypes at the time. Unless otherwise specified, each buffer was freshly supplemented with 70 mM 2-Mercaptoethanol and completeM Protease Inhibitor Cocktail (Sigma). For each genotype, we harvested on ice about 5 ml of unopened flower buds including one open flower for each inflorescence. The spores were released from the buds in a prechilled mortar on ice, using 5 ml of 0.1 M mannitol, by gently tapping with a prechilled pestle for 1-2 minutes. The suspension was transferred to a 50 ml conical tube, and an additional 10 ml of 0.1 M mannitol were used to rinse the mortar. The tube was then vortexed intermittently for 30 seconds to release the spores, and the suspension was filtered through a 100 µm nylon membrane to remove the tissue. An additional 5 ml of 0.1 M mannitol were used to rinse the tube and poured over the same filter, obtaining a total of 20 ml of suspension for each sample. The spores were further filtered twice through a 60 µm nylon mesh and then divided into two 15-ml glass tubes. The tubes were spun with a Sorvall Lynx 4000 Centrifuge (Thermo Scientific) with TH13-6x50 swing-out rotor, for 10 min at 900 g, 4°C, acceleration speed 5, braking speed 8. The supernatant was carefully removed, and the pellets were resuspended in 1 ml of ice-cold 0.1 M mannitol. Each aliquot was transferred to a new tube and layered above 3 ml of 20% Percoll (diluted in 0.1 M Mannitol). The samples were centrifuged at 450 g for 10 min at 4°C, with acceleration speed 5 and braking speed 8. The two pellets from the same genotype were combined with 2 ml of 0.1 M mannitol, and the centrifugation over 20% Percoll was repeated two times to clean the spores further. The purified mixed spores were then transferred to a 1.5 ml tube and inspected under the microscope.

The protocol for nuclei extraction and purification was adapted from (Santos et al., 2017). The spores were pelleted via 5 min centrifugation at 500 g, 4°C. The pellets were resuspended in 800 μl of Galbraith buffer supplemented with 70 mM 2-Mercaptoethanol (45 mM MgCl2, 30 mM C6H5Na3O7.2H2O [Trisodium citrate dihydrate], 20 mM MOPS, 0.1% [v/v] Triton X-100, pH 7). The suspension was transferred to a 1.5 ml tube containing 100 μl of acid-washed 0.5 mm glass beads (Sigma). To break the pollen walls, the samples were vortexed at max speed for 2 min total in a cold room with the following vortexing protocol: 7 sec vortex, 3 sec invert for the first minute followed by 7 sec vortex, 2 sec invert for the second minute. We then filtered the nuclei by briefly spinning them over a 10 μm cellTrics filter placed in a clean 1.5 ml tube. 400 μl of Galbraith buffer were used to rinse the beads, and added to the cellTrics filter. While keeping the flowthrough on ice, the unbroken pollen grains that remained on the filter were collected by pipetting with 800 μl of Galbraith buffer, and were placed back in the tube with the glass beads. An additional round of vortexing and filtering was performed as before, to increase the nuclei yield. The nuclei in the two tubes were then spun down for 5 min at 500 g, 4°C, and the pellets were resuspended by gentle pipetting with 50 μl of CyStain UV Precise P - Nuclei Extraction Buffer (Sysmex, 05-5002-P02). To stain the nuclei we then added 400 μl of CyStain UV Precise P - Staining Buffer (Sysmex, 05-5002-P01) and we supplemented the sample with Protector RNase Inhibitor (Sigma) to a final concentration of 0.2 U/µl. The samples were passed over the filter of a FACS tube (Falcon 352235) and immediately sorted. Sorting was done with a BD FACS ARIAII instrument equipped with a 355nm UV laser, using the 70 μm nozzle. The gating strategy is shown in Figure S5A-C. For each sample, we sorted 40,000-60,000 nuclei in 500 µl of Nuclei wash buffer (2% BSA in 1X PBS) supplemented with Protector RNase Inhibitor (Sigma) to a final concentration of 0.2 U/µl. The sorted nuclei were pelleted by centrifugation for 5 min at 500 g, 4°C. The pellet was resuspended in 20-25 μl of buffer and the entire sample was used as input for the 10x Genomics Chromium Single Cell 3’ Reagent Kit v3. The subsequent steps were performed according to the manufacturer’s instructions.

### QUANTIFICATION AND STATISTICAL ANALYSIS

#### Analysis of BS-seq

The BS-seq data present in this paper was reanalyzed from the following deposited datasets. Wild-type flower BS-seq: GSM5026060 and GSM5026061 merged replicates (Ichino et al., 2021). Wild-type sorted SN: GSM952445 (Ibarra et al., 2012). Wild-type sorted VN: GSM952447 (Ibarra et al., 2012). Raw reads were trimmed with TrimGalore (Babraham Institute) and mapped to the TAIR10 genome with Bismark (Krueger and Andrews, 2011). Bismark was also used to obtain the methylation percentages for each cytosine and to generate the per-position DNA methylation tracks. The quantification of the average methylation percentage at promoters was calculated with bedtoolsmap (Quinlan and Hall, 2010) with the option “*mean*”. Promoters were defined as a 600 bp region surrounding the TSS.

#### Analysis of bulk RNA-seq

The bulk RNA-seq data was analyzed as previously described (Ichino et al., 2021). The RNA-seq reads were filtered based on quality score and trimmed to remove Illumina adapters using Trim Galore (Babraham Institute). The filtered reads were mapped to the Arabidopsis reference genome (TAIR10) using STAR (Dobin et al., 2013), allowing 5% of mismatches (-outFilterMismatchNoverReadLmax 0.05) and unique mapping (-- outFilterMultimapNmax 1). PCR duplicates were removed using MarkDuplicates from the Picard Tools suite. Coverage tracks for visualization in the genome browser were generated using Deeptools 3.0.2 bamCoverage with the options --normalizeUsing RPKM and --binSize 10 (Ramírez et al., 2016). The number of reads mapping to genes or transposable elements were determined using HTseq (Anders et al., 2015) with the option --mode=union. We used as reference the transcriptome annotations generated as described in the paragraph “Pollen transcriptome reannotation”, and all transcripts were analyzed together in the DEG analysis (genes, TEs, and other undefined non-coding transcripts). The HTseq gene counts were used to perform the differential gene expression analysis using the R package DEseq2 (Love et al., 2014) with a cutoff for significance of padj <0.05 and |log2FC|>0.5. The transcripts per million (TPM) values were estimated using Kallisto version 0.46.0 (Bray et al., 2016). Figures were generated using the R package *ggplot*. The heatmaps of RNA-seq and methylation data were made with the R package *ComplexHeatmap* (Gu et al., 2016).

The TE family analysis was done using the package *TEtranscripts* (Jin et al., 2015). Reads were mapped with STAR allowing multimapping up to 100 hits (-- outFilterMultimapNmax 100 and -winAnchorMultimapNmax 100). *TEtranscripts* was run with default options (--mode multi). Significantly upregulated TE families were defined as the ones with adjusted p-value < 0.05.

The gene ontology (GO) analysis of the *met1* DEGs was done with the R package *clusterProfiler* (Wu et al., 2021), running *enrichGO* with the following parameters: ont = “all”, pvalueCutoff = 0.05, qvalueCutoff = 0.10. We used as reference genes the list of all genes with baseMean>2 (from the DEseq2 table).

The following RNA-seq datasets were downloaded from GEO and reanalyzed as described above: wild-type VN (GSM4700179, GSM4700180, GSM4700181) (Borg et al., 2021a), wild-type sperm (GSM4700188, GSM4700189, GSM4700190) (Borg et al., 2020), wild-type and *mbd5/6* flower buds (GSM5026083 to GSM5026091) (Ichino et al., 2021), wild-type, *sln* and *met1* flower buds (GSM5026092 to GSM5026094 and GSM5026098 to GSM5026103) (Ichino et al., 2021), wild-type and *met1* seedlings (GSM938342, GSM938343, GSM938348, GSM938349) (Stroud et al., 2012).

The curated list of “pollen-tube related genes” that are demethylated by DME, used in Figure 6B, includes: AT1G66235, AT2G19480, AT1G44120, AT2G16586, AT2G14260, AT5G28470, AT1G35540, AT3G30720, AT1G74800, AT2G16015, AT2G22055/RALFL15, AT2G07040/PRK2, novel_Chr1_coding_102/RKF2/AT1G19090, novel_Chr2_coding_224/PLOU/AT2G16030.

#### Pollen transcriptome reannotation

All Pollen RNA-seq libraries (27 libraries total) were filtered and trimmed as described in previous section (“Analysis of bulk RNA-seq”), and aligned to the Arabidopsis reference genome (TAIR10) using STAR (Dobin et al., 2013). STAR genome indexes were made using the Araport11 annotations from Cheng et al., 2017. BAM files for the 27 pollen RNA-seq samples were pooled into a single BAM file using samtools merge, with a total of approximately 450M reads in the pool. Note all libraries were made using the Truseq stranded RNA kit and shared the same ’strandedness’. Transcripts were assembled from this BAM file using four different programs: Trinity v2.13.2 (Haas et al., 2013), Cufflinks v.2.2.1 (Trapnell et al., 2010), CLASS2 v.2.1.7 (Song et al., 2016) and StringTie v.2.1.6 (Pertea et al., 2015). When annotation information could also be provided to guide assembly (Trinity, Cufflinks, and StringTie), Araport11 was again provided. Trinity was run with options `--genome_guided_bam pooled.bam -- genome_guided_max_intron 2000 --jaccard_clip --SS_lib_type RF`. The resulting fasta file was converted to GTF using gmap version 2021-08-25 (Wu and Watanabe, 2005). Cufflinks was run with options `-g araport11.gtf -I 5000 -- library-type fr-firststrand --max-bundle-length 30000 --min-intron-length 15 --overlap-radius 1`. CLASS2 was run with default options, and StringTie with options `-G araport11.gtf --rf -m 40 -g 1`. Additionally, junctions were detected using Portcullis v.1.2.2 (Mapleson et al., 2018) with options `-- orientation FR --strandedness firststrand --max_length 2000`. Assembled transcripts from each of these sources were combined and best transcripts were selected using Mikado v.2.3.2 (Venturini et al., 2018). Mikado is run in 4 parts: (1) configure with options `--strand-specific` and providing Portcullis predictions to `-- junctions`, (2) prepare with default options; this step pools predicted transcripts from all input sources (3) serialise with `--orfs` = ORFs predicted by prodigal v.2.6.3 (Hyatt et al., 2010) with options `-g1 -f gff` and `-xml` blastx output from BLASTX v.2.11.0 with options `-max_target_seqs 5 -outfmt “6 qseqid sseqid pident length mismatch gapopen qstart qend sstart send evalue bitscore ppos btop”`, and (4) pick, with options `--scoring-file plant.yaml --no-purge` (plant.yaml from Mikado Github). Mikado selections were further refined using custom python script mikado_refine.py (available from Github at https://github.com/clp90/mbd56_pollen), which incorporates additional RNA-seq coverage information and a few other selection parameters, with default options. To simplify our analysis, mikado_refine.py outputs a single ’best’ representative annotation for each gene. Final updated annotations and codes used to generate them are available from Github at https://github.com/clp90/mbd56_pollen.

#### Analysis of ATAC-seq

The bigWig track files for the wild-type VN, GN, and SN ATAC-seq datasets were downloaded from GEO (GSM4699541 to GSM4699544, and GSM5027098 GSM5027100) (Borg et al., 2021a). The ATAC-seq enrichment at promoters was obtained with deeptools multiBigwigSummary with the -BED option and --outRawCounts (Ramírez et al., 2016). The promoter regions were defined as windows from 300 bp before the TSS until 300 bp after the TSS.

#### Analysis of snRNA-seq

##### Preprocessing

Cell Ranger 6.1.1 software (10X genomics) was used to process the raw data. The reads were aligned to the Arabidopsis reference annotations generated as explained in “Pollen transcriptome reannotations”. For each individual sample, we then removed the ambient RNA using SoupX with standard settings (Young and Behjati, 2020). The data was then imported in Seurat 4.0.4 (Hao et al., 2021) removing cells in which less than 300 genes were detected. The data was normalized with *NormalizeData* (normalization.method = “LogNormalize”, scale.factor *= 10000*), and scaled with *ScaleData* with default settings. We then performed principal component analysis on all genes with *RunPCA* (npcs=20) and dimensionality reduction with *RunUMAP* using the first 20 principal components (PCs). Next, we used DoubletFinder v3 (McGinnis et al., 2019) to identify doublets with the standard workflow based on the first 20 principal components, and we used *find.pK* to identify the optimal pK parameter for each sample. The pK values and the percentage of doublets removed for each sample are available in Table S3. Lastly, we removed nuclei with more than 5% of mitochondrial reads or more than 15% of chloroplast reads.

##### Integration and clustering

The different datasets were integrated with Seurat v4 *FindIntegrationAnchors* and *IntegrateData* using default settings (nfeatures = 2000). The integrated data was scaled with *ScaleData* with default settings, and PCA analysis was performed with *RunPCA* (npcs=40). Dimensionality reduction was done with *RunUMAP* using the first 40 PCs. We then performed clustering analysis using *FindNeighbors* to calculate the k-nearest neighbors based on the first 40 PCs, and *FindClusters* with resolution = 0.3, using the Louvain algorithm (default). The list of numbers of cells per cluster for each sample is available in Table S3. We note that the relative number of cells per cluster was quite variable when comparing different wild-type datasets, likely because of the variability introduced while harvesting the inflorescences. Therefore, we think that this experimental approach cannot be used to determine how a given mutation can impact the speed of specific stages of pollen development, unless the phenotype is very strong such as in the case of *met1* (Figure 5A).

##### Cluster markers and gene ontology (GO) analysis

The cluster markers (available in Table S4) were obtained with the Seurat function *FindAllMarkers* ran on the Col0 samples only, using the options only.pos = TRUE and logfc.threshold = 0.1. This generated a non-stringent list of markers that were ranked by average log2FC. To perform the GO analysis, we selected the markers with p_val_adj < 0.05 and avg_log2FC > 1, and we used as reference the list of all Arabidopsis genes included in our reference transcriptome. The analysis was done with the R package *clusterProfiler* (Wu et al., 2021), running *enrichGO* with the following parameters: ont = “all”, pvalueCutoff = 0.05, qvalueCutoff = 0.10. The complete list of results is available in Table S6.

##### Pseudotime analysis

The VN pseudotime analysis was performed with Monocle3 (Cao et al., 2019; Qiu et al., 2017; Trapnell et al., 2014) following the standard workflows. The Seurat object was subsetted to select the Col0 cells assigned to the VN clusters. The data was then converted into a Monocle3 cell_data_set object with the R package “*SeuratWrappers*”. The trajectory was constructed with the *learn_graph* function and the root was manually selected with the *order_cells* function (reduction_method = “UMAP”). To find the genes that vary as a function of pseudotime, we used the *graph_test* function with the option neighbor_graph=“principal_graph”, to test whether cells at similar positions on the trajectory have correlated expression. We then selected the genes with q_value < 0.01 & morans_I > 0.1. The Seurat function *AverageExpression* with the option slot = “data” was used to obtain the average expression levels of those genes (log normalized), for all the cells belonging to a given pseudotime interval (binning of the pseudotime in 43 intervals). The average expression values were scaled by z-score, visualized in a heatmap and clustered with the R package *pheatmap* (using *hclust*). Gene ontology analysis was then performed on each group of genes as described in the paragraph “Cluster markers and gene ontology (Go) analysis”.

##### Differential gene expression (DEG) analysis of snRNA-seq data

Given the well described challenges of performing DEG analysis with single cell RNA-seq data, we tested several different approaches, including both pseudobulk and single cell methods as suggested in the literature (Squair et al., 2021). The pseudobulk methods that take into account biological replicates are considered more robust; however, when we used either pseudobulk or single cell methods to identify the *mbd5/6* DEGs, we observed a large number of DEGs that corresponded to highly expressed and unmethylated genes, and therefore don’t have the typical features of MBD5/6 targets. Thus, we decided to take advantage of the 6 independent wild-type replicates that we generated on different days to compile a list of “noise DEGs” including genes that are called as differentially expressed when comparing wild-types with each other. We then subtracted these genes from the lists of DEGs obtained for each mutant compared to its matched wild-type (Figure S6A,B). Without filtering the “noise DEGs”, most of the *mbd5/6* up-DEGs are unmethylated genes that are not upregulated in mature pollen bulk RNA-seq (Figure S6C), while after filtering, almost all the DEGs have promoter methylation and are upregulated in bulk RNA-seq as well (Figure 4C). This observation gave us confidence in the validity of our method. We do however acknowledge that this approach is very stringent and could bias the analysis towards genes that are lowly or not expressed in wild-type. For the purpose of this study however this is not a concern because we are investigating derepression mutants, which mostly affect loci that are lowly or not expressed in wild-type. In the next paragraph we explain in more detail the methods used to analyze and plot the data.

The DEG analysis was done by running on each individual cluster the Seurat function *FindMarkers* with “wilcox” as statistical method. All transcripts were considered together for the DEG analysis (genes, TEs, and other non-coding transcripts). Each mutant was compared to its matched wild-type control (processed on the same day). We also compared all the wild-type samples to each other, and we selected the significant DEGs (p_val_adj<0.05) to obtain the list of “noise DEGs”, which were filtered out of each list of mutant DEGs. Downstream analyses were done using the filtered mutant DEGs that had an avg_log2FC < -0.25 or > 0.25.

To visualize the data, we grouped together nuclei clusters that were similar. The cluster groups were defined as follows: MN1, MN2, and MN3 were combined into the “MN” group, VN1, VN2 and VN3 were combined into the “early VN” group, VN4 and VN5 in the “late VN” group, GN1 and GN2 in the “GN” group. The heatmaps were generated by first extracting from the Seurat object the average expression values per cluster group, using the Seurat function *AverageExpression* with the option slot = “data”. These values were then scaled by calculating for each gene the z-score across all columns. To combine the snRNA-seq, BS-seq, and bulk RNA-seq in a heatmap we used the R package *ComplexHeatmap*. The boxplots of log2FC were generated with the R package *ggplot*, using the average log2FC values for each cluster group obtained using the Seurat function *FindMarkers*.

### ADDITIONAL RESOURCES

The snRNA-seq can be inspected with an interactive website:

https://singlecell.mcdb.ucla.edu/snRNAseq_pollen/

### SUPPLEMENTAL ITEMS

Table S1 all genomics datasets-Related to Figures 1,2,6,7.xlsx

Table S2 bulk pollen DEGs - Related to Figure 2.xlsx

Table S3 cellranger and DoubletFinder - Related to Figure 3.xlsx Table S4 cluster markers - Related to Figure 3.csv

Table S5 GO VN lineage - Related to Figure 4.xlsx

Table S6 GO all clusters - Related to Figure 3.xlsx

Table S7 10X DEGs - Related to Figure 4 and 5.xlsx

## KEY RESOURCES TABLE

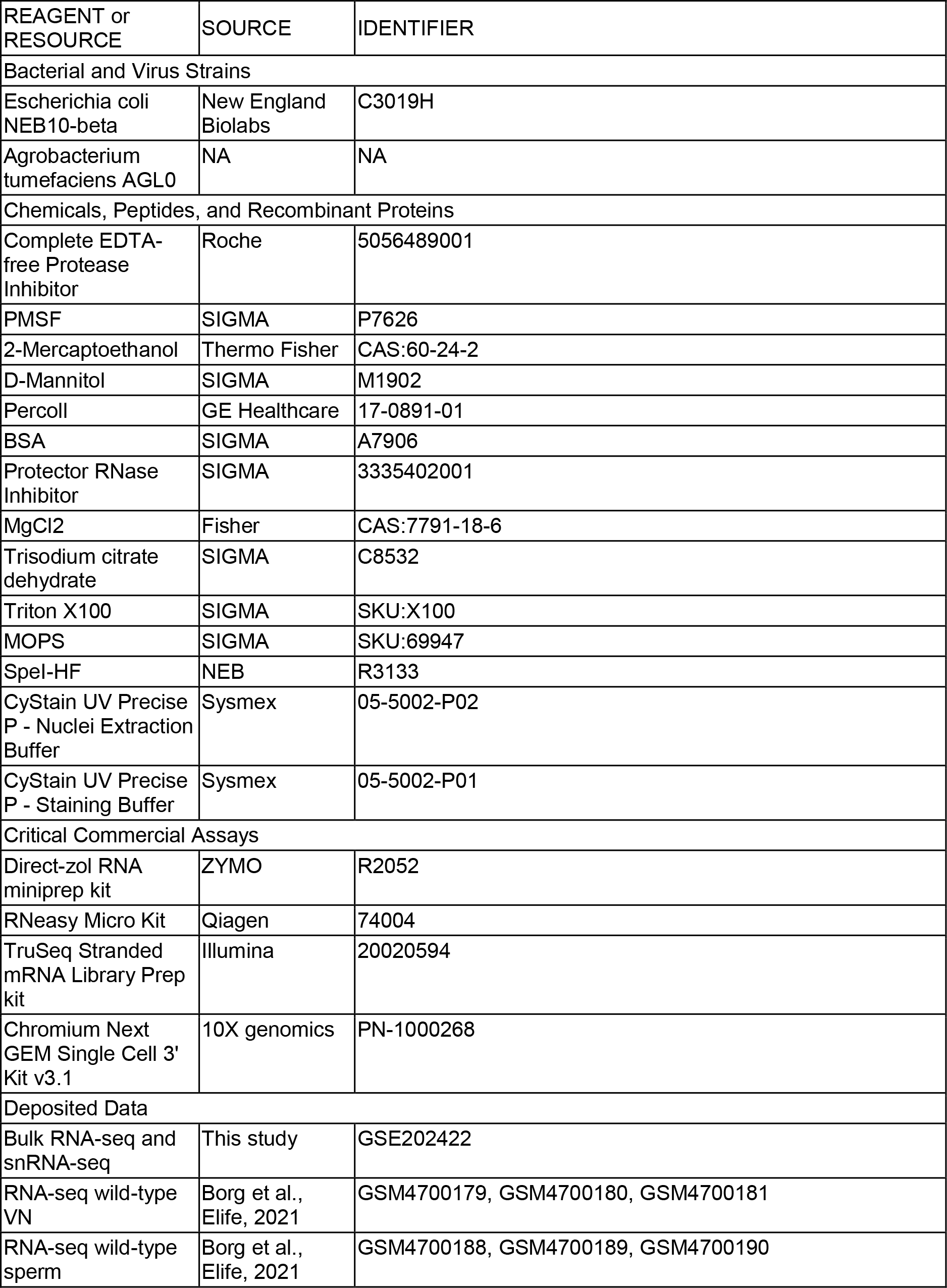
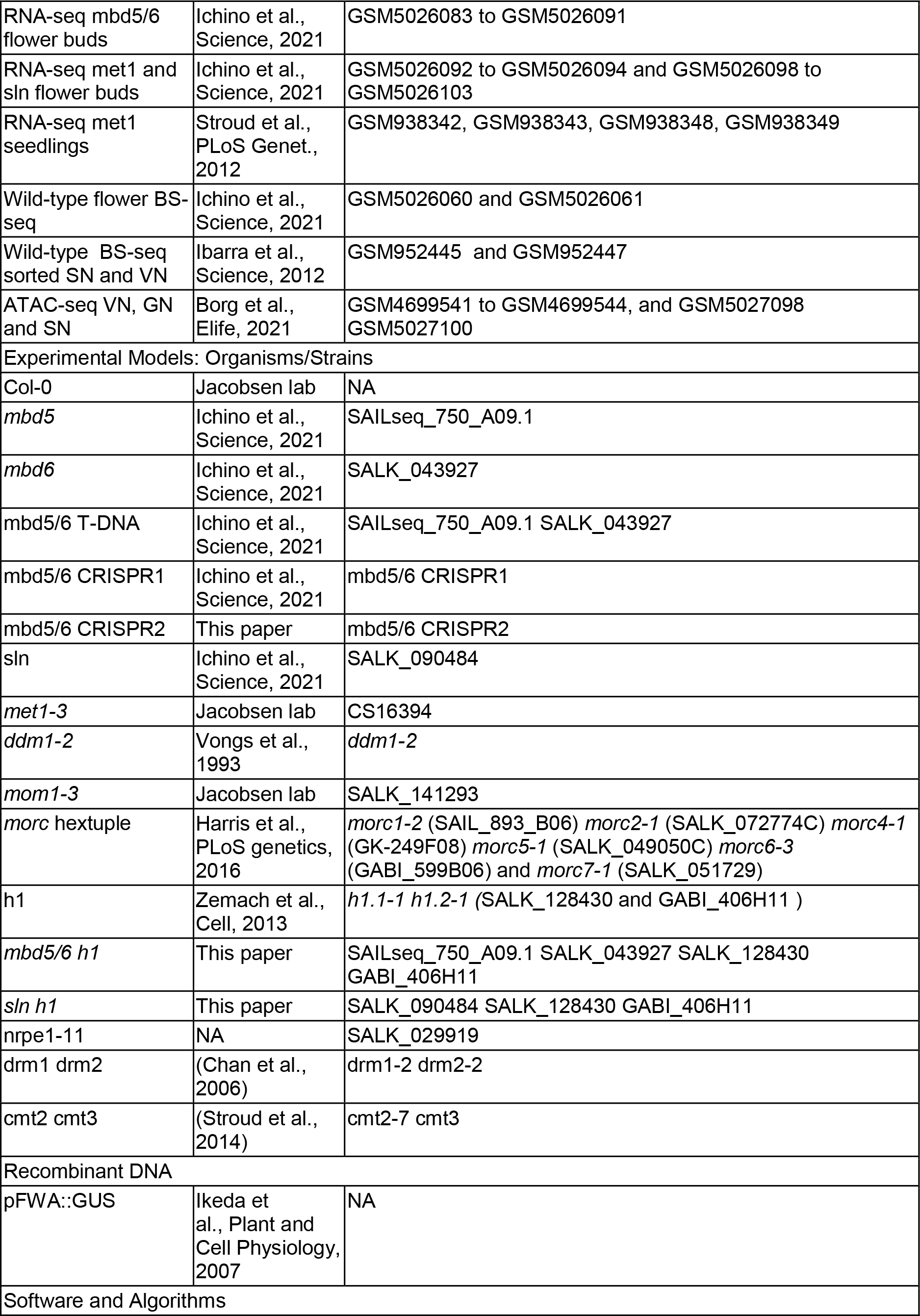
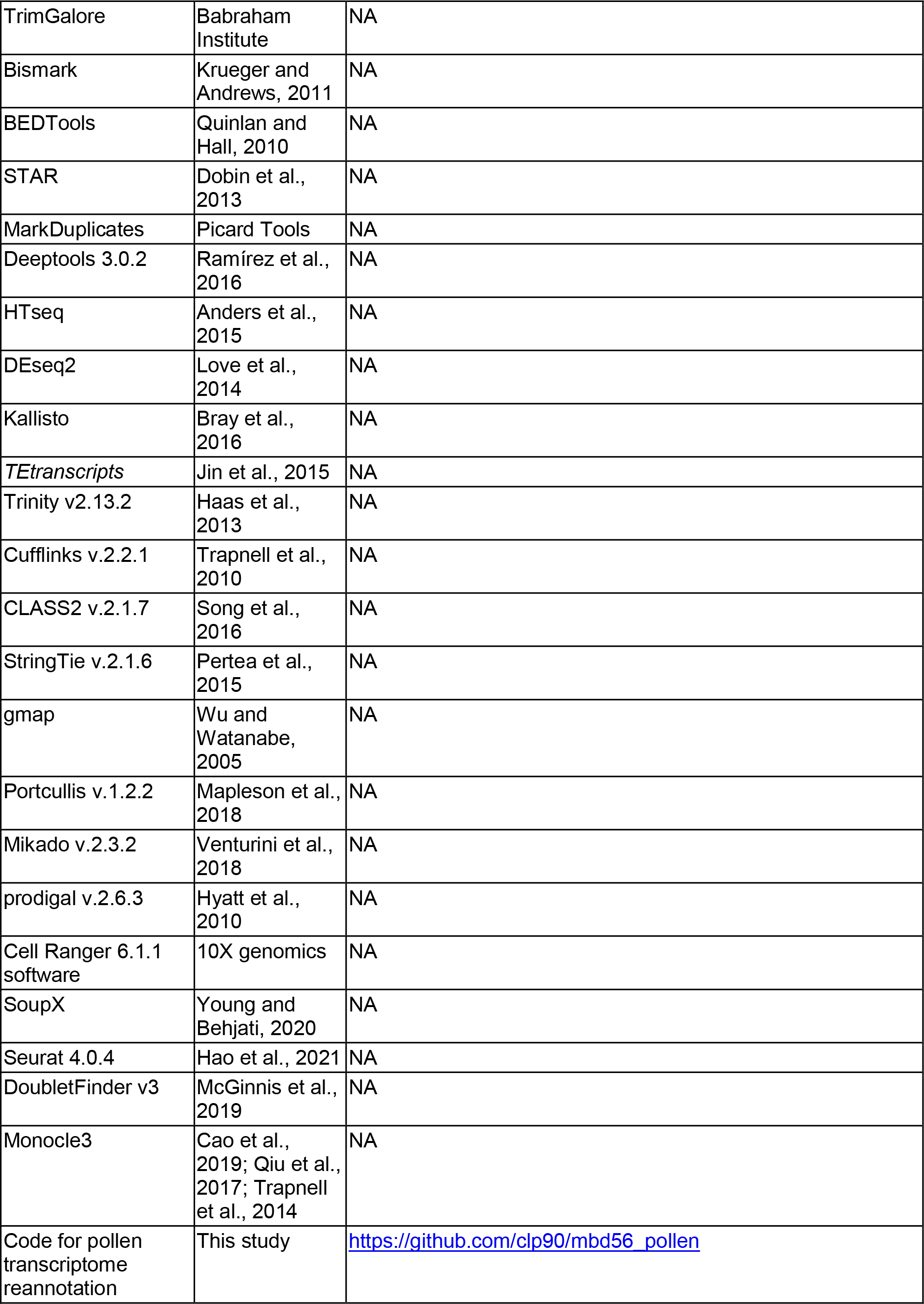

| REAGENT or RESOURCE | SOURCE | IDENTIFIER |
| --- | --- | --- |
| **Bacterial and Virus Strains** |  |  |
| Escherichia coli NEB10-beta | New England Biolabs | C3019H |
| Agrobacterium tumefaciens AGL0 | NA | NA |
| **Chemicals, Peptides, and Recombinant Proteins** |  |  |
| Complete EDTA-free Protease Inhibitor | Roche | 5056489001 |
| PMSF | SIGMA | P7626 |
| 2-Mercaptoethanol | Thermo Fisher | CAS:60-24-2 |
| D-Mannitol | SIGMA | M1902 |
| Percoll | GE Healthcare | 17-0891-01 |
| BSA | SIGMA | A7906 |
| Protector RNase Inhibitor | SIGMA | 3335402001 |
| MgCl<sub>2</sub> | Fisher | CAS:7791-18-6 |
| Trisodium citrate dehydrate | SIGMA | C8532 |
| Triton X100 | SIGMA | SKU:X100 |
| MOPS | SIGMA | SKU:69947 |
| SpeI-HF | NEB | R3133 |
| CyStain UV Precise P - Nuclei Extraction Buffer | Sysmex | 05-5002-P02 |
| CyStain UV Precise P - Staining Buffer | Sysmex | 05-5002-P01 |
| **Critical Commercial Assays** |  |  |
| Direct-zol RNA miniprep kit | ZYMO | R2052 |
| RNeasy Micro Kit | Qiagen | 74004 |
| TruSeq Stranded mRNA Library Prep kit | Illumina | 20020594 |
| Chromium Next GEM Single Cell 3' Kit v3.1 | 10X genomics | PN-1000268 |
| **Deposited Data** |  |  |
| Bulk RNA-seq and snRNA-seq | This study | GSE202422 |
| RNA-seq wild-type VN | Borg et al., Elife, 2021 | GSM4700179, GSM4700180, GSM4700181 |
| RNA-seq wild-type sperm | Borg et al., Elife, 2021 | GSM4700188, GSM4700189, GSM4700190 |
| RNA-seq mbd5/6 flower buds | Ichino et al., Science, 2021 | GSM5026083 to GSM5026091 |
| RNA-seq met1 and sln flower buds | Ichino et al., Science, 2021 | GSM5026092 to GSM5026094 and GSM5026098 to GSM5026103 |
| RNA-seq met1 seedlings | Stroud et al., PLoS Genet., 2012 | GSM938342, GSM938343, GSM938348, GSM938349 |
| Wild-type flower BS-seq | Ichino et al., Science, 2021 | GSM5026060 and GSM5026061 |
| Wild-type BS-seq sorted SN and VN | Ibarra et al., Science, 2012 | GSM952445 and GSM952447 |
| ATAC-seq VN, GN and SN | Borg et al., Elife, 2021 | GSM4699541 to GSM4699544, and GSM5027098 GSM5027100 |
| Experimental Models: Organisms/Strains |  |  |
| Col-0 | Jacobsen lab | NA |
| *mbd5* | Ichino et al., Science, 2021 | SAILseq_750_A09.1 |
| *mbd6* | Ichino et al., Science, 2021 | SALK_043927 |
| *mbd5/6* T-DNA | Ichino et al., Science, 2021 | SAILseq_750_A09.1 SALK_043927 |
| *mbd5/6* CRISPR1 | Ichino et al., Science, 2021 | *mbd5/6* CRISPR1 |
| *mbd5/6* CRISPR2 | This paper | *mbd5/6* CRISPR2 |
| *sln* | Ichino et al., Science, 2021 | SALK_090484 |
| *met1-3* | Jacobsen lab | CS16394 |
| *ddm1-2* | Vongs et al., 1993 | *ddm1-2* |
| *mom1-3* | Jacobsen lab | SALK_141293 |
| *morc* hextuple | Harris et al., PLoS genetics, 2016 | *morc1-2* (SAIL_893_B06) *morc2-1* (SALK_072774C) *morc4-1* (GK-249F08) *morc5-1* (SALK_049050C) *morc6-3* (GABI_599B06) and *morc7-1* (SALK_051729) |
| *h1* | Zemach et al., Cell, 2013 | *h1.1-1 h1.2-1* (SALK_128430 and GABI_406H11 ) |
| *mbd5/6 h1* | This paper | SAILseq_750_A09.1 SALK_043927 SALK_128430 GABI_406H11 |
| *sln h1* | This paper | SALK_090484 SALK_128430 GABI_406H11 |
| *nrpe1-11* | NA | SALK_029919 |
| *drm1 drm2* | (Chan et al., 2006) | *drm1-2 drm2-2* |
| *cmt2 cmt3* | (Stroud et al., 2014) | *cmt2-7 cmt3* |
| Recombinant DNA |  |  |
| pFWA::GUS | Ikeda et al., Plant and Cell Physiology, 2007 | NA |
| Software and Algorithms |  |  |
| TrimGalore | Babraham Institute | NA |
| Bismark | Krueger and Andrews, 2011 | NA |
| BEDTools | Quinlan and Hall, 2010 | NA |
| STAR | Dobin et al., 2013 | NA |
| MarkDuplicates | Picard Tools | NA |
| Deeptools 3.0.2 | Ramírez et al., 2016 | NA |
| HTseq | Anders et al., 2015 | NA |
| DEseq2 | Love et al., 2014 | NA |
| Kallisto | Bray et al., 2016 | NA |
| *TEtranscripts* | Jin et al., 2015 | NA |
| Trinity v2.13.2 | Haas et al., 2013 | NA |
| Cufflinks v.2.2.1 | Trapnell et al., 2010 | NA |
| CLASS2 v.2.1.7 | Song et al., 2016 | NA |
| StringTie v.2.1.6 | Pertea et al., 2015 | NA |
| gmap | Wu and Watanabe, 2005 | NA |
| Portcullis v.1.2.2 | Mapleson et al., 2018 | NA |
| Mikado v.2.3.2 | Venturini et al., 2018 | NA |
| prodigal v.2.6.3 | Hyatt et al., 2010 | NA |
| Cell Ranger 6.1.1 software | 10X genomics | NA |
| SoupX | Young and Behjati, 2020 | NA |
| Seurat 4.0.4 | Hao et al., 2021 | NA |
| DoubletFinder v3 | McGinnis et al., 2019 | NA |
| Monocle3 | Cao et al., 2019; Qiu et al., 2017; Trapnell et al., 2014 | NA |
| Code for pollen transcriptome reannotation | This study | https://github.com/clp90/mbd56_pollen |

## Supplementary figures

**Figure S1 – related to Figure 1:**
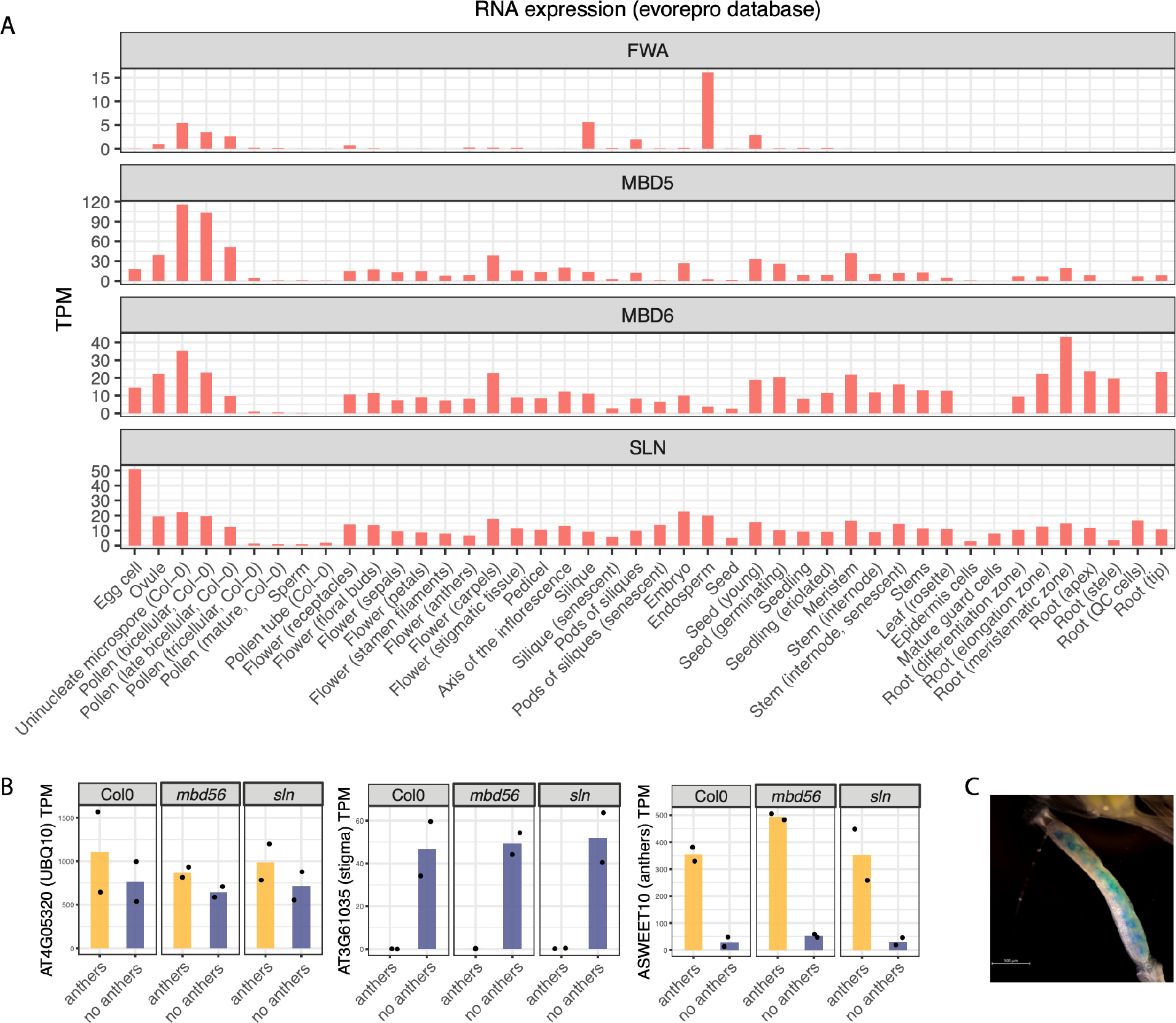
Tissue specificity of *FWA* expression. A) Barplots of average expression for the indicated genes (transcripts per million [TPM]) across a panel of tissues. The data was downloaded from https://evorepro.sbs.ntu.edu.sg and re-plotted. B) RNA-seq data of dissected flower buds (dissection strategy shown in Figure 1A). Shown are the average TPM values for a constitutively expressed gene (*POLYUBIQUITIN 10*), a gene enriched in stigmatic tissue (AT3G61035), and an anthers marker (*ASWEET10*). The dots indicate individual replicates. C) Representative image of the endogenous expression of *FWA* in early developing seeds. The *pFWA::GUS* reporter line was GUS-stained and cleared with ClearSee (Kurihara et al., 2015). Scale bar: 500 μm.

**Figure S2 – related to Figure 2:**
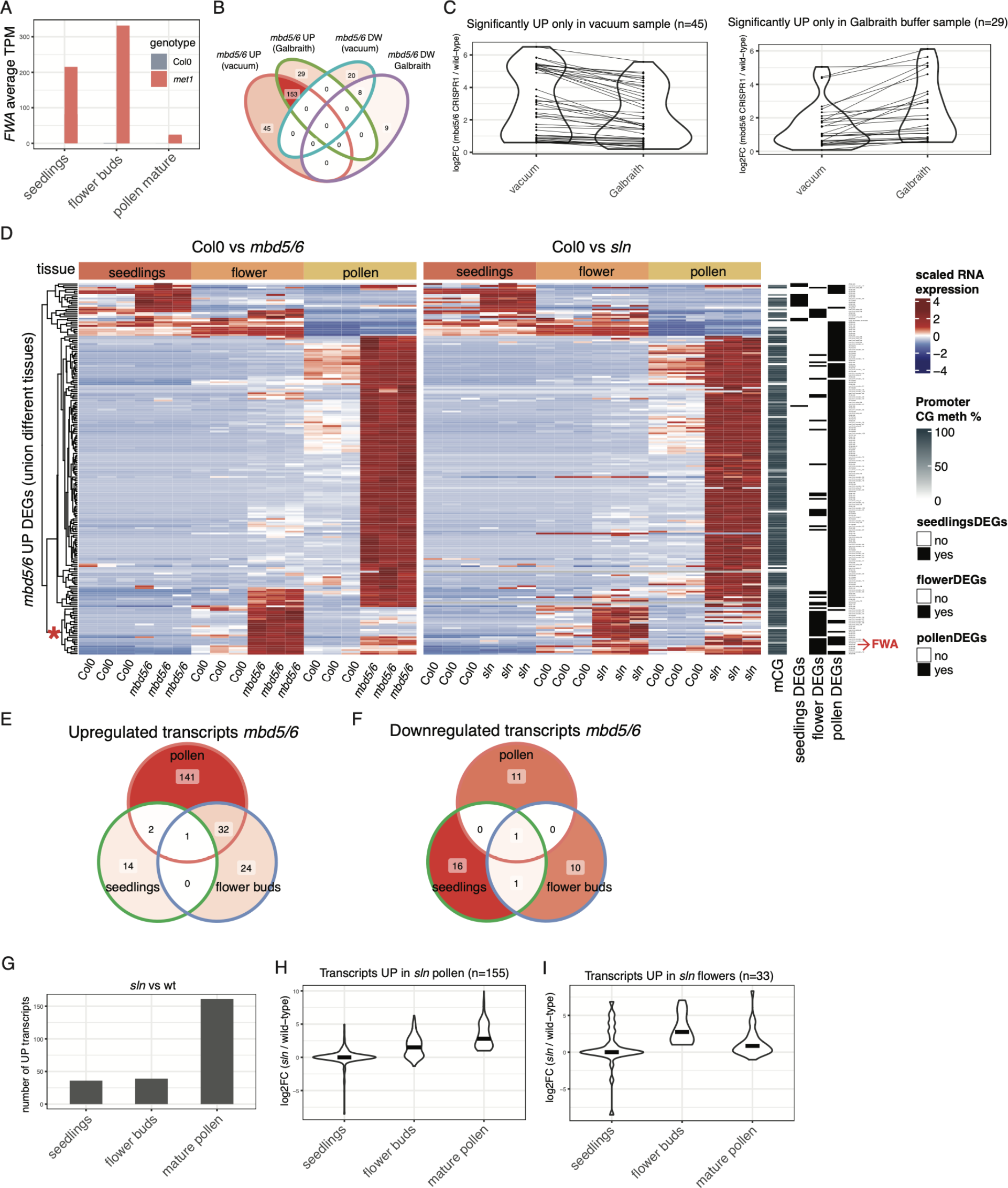
Tissue specificity of the *mbd5/6* and *sln* transcriptional derepression phenotypes. A) *FWA* average expression in the indicated tissues measured by RNA-seq. The *met1* seedlings data is from (Stroud et al., 2012) and the flower buds data is from (Ichino et al., 2021). B) Venn diagram of overlaps between the upregulated DEGs detected in *mbd5/6* pollen extracted with two different methods: vacuum aspiration or Galbraith buffer (see Methods). C) Violin plots showing the differential expression level (log2FC) of the transcripts that were called as significantly upregulated in only one of the two methods (vacuum aspiration or Galbraith buffer). D) Heatmap showing the union of the *mbd5/6* upregulated transcripts in seedlings, flower buds and mature pollen (n=204). The RNA-seq data is shown as z-score of TPM, the methylation data is the average CG methylation percentage in flower buds, in a 600 bp window centered on the TSS. The row annotations on the right indicate whether each gene was called as significant *mbd5/6* DEG in each tissue. The red asterisk on the dendrogram indicates a group of genes that are more strongly upregulated in flowers than in pollen. This group includes *FWA*, as shown on the right. E,F) Venn diagrams of upregulated and downregulated transcripts in *mbd5/6* seedlings, flower buds, and mature pollen. G) Number of upregulated transcripts in *sln* vs Col0 (wild-type) in the indicated tissues. The *sln* flower buds RNA-seq is from (Ichino et al., 2021). H,I) Distribution of the *sln* vs Col0 log2 fold-changes for the indicated transcripts in different tissues. Black bar indicates the median.

**Figure S3 – related to Figure 2:**
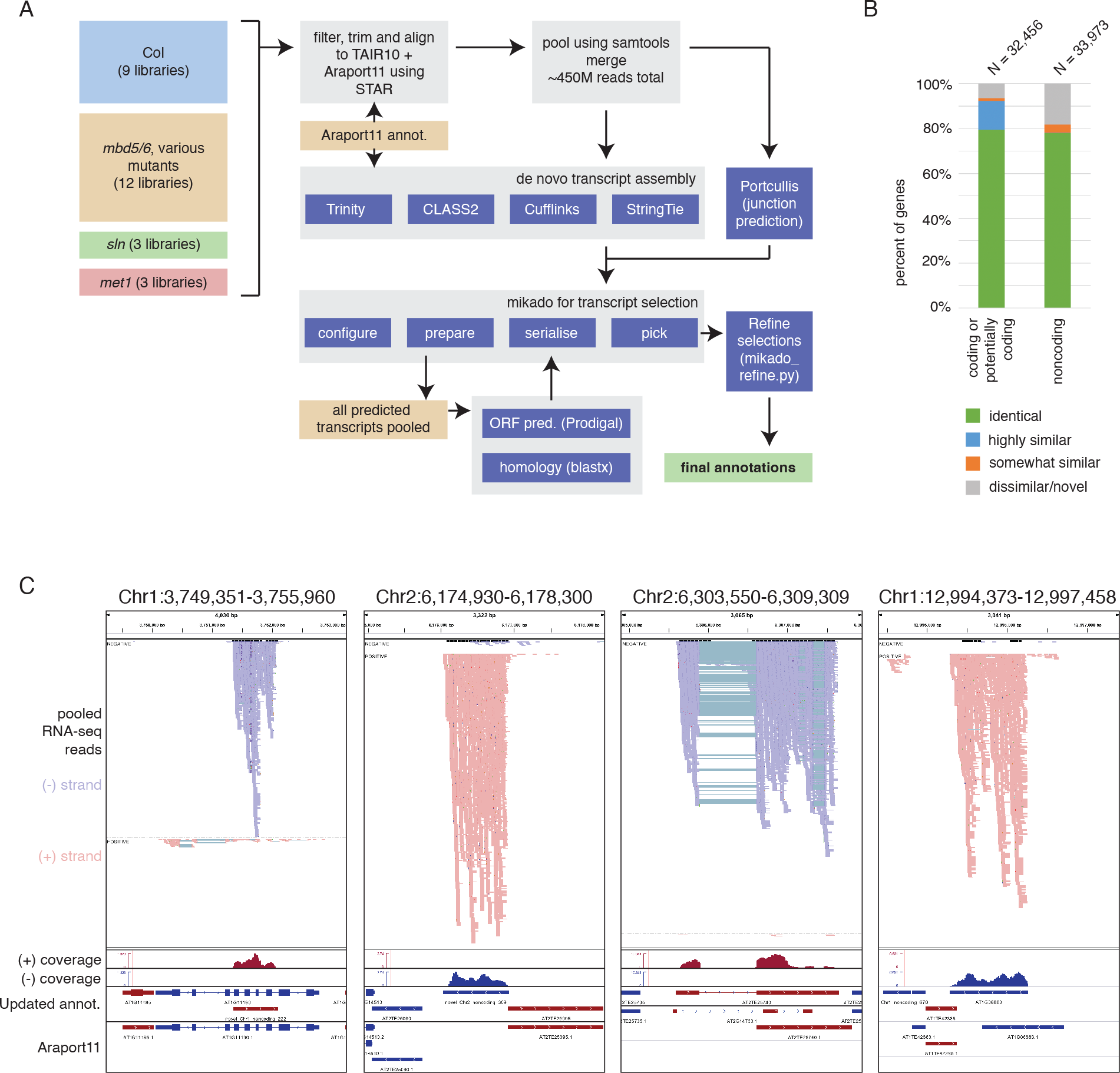
Reannotation of *A. thaliana* transcriptome based on pollen data from *mbd5/6*, *sln*, and *met1*. A) Overview of approach used to obtain reannotation, using a combination of four transcript assembly programs: Trinity v2.13.2 (Haas et al., 2013), Cufflinks v.2.2.1 (Trapnell et al., 2010), CLASS2 v.2.1.7 (Song et al., 2016) and StringTie v.2.1.6 (Pertea et al., 2015). The program Mikado was used to select the best transcript. Transcript selection was further refined using a custom script (available on Github). B) Percentage of genes that were identical, highly similar, somewhat similar, and dissimilar/novel in the reannotation vs. in the original Araport11 annotations. Identical = all transcript features unchanged, highly similar (coding) = CDS unchanged but altered UTRs, highly similar (noncoding) = > 95% overlap of exons, somewhat similar (coding) = CDS > 80% similar and in frame, somewhat similar (noncoding) = > 50% overlap of exons, all others considered dissimilar or novel. C) Example loci showing missing annotations added during reannotation (left two panels), and example loci where existing annotations were improved (right two panels). Top track shows reads from pooled BAM file, next two tracks indicate coverage originating from (+) or (-) strand. Top annotation track (“Updated annot.” shows updated annotations while starting annotations from Araport11 are shown at bottom. Forward annotations are colored red, reverse colored blue. Note that reads mapping to (-) strand originate from (+) strand annotations and vice versa.

**Figure S4 – related to Figure 2.**
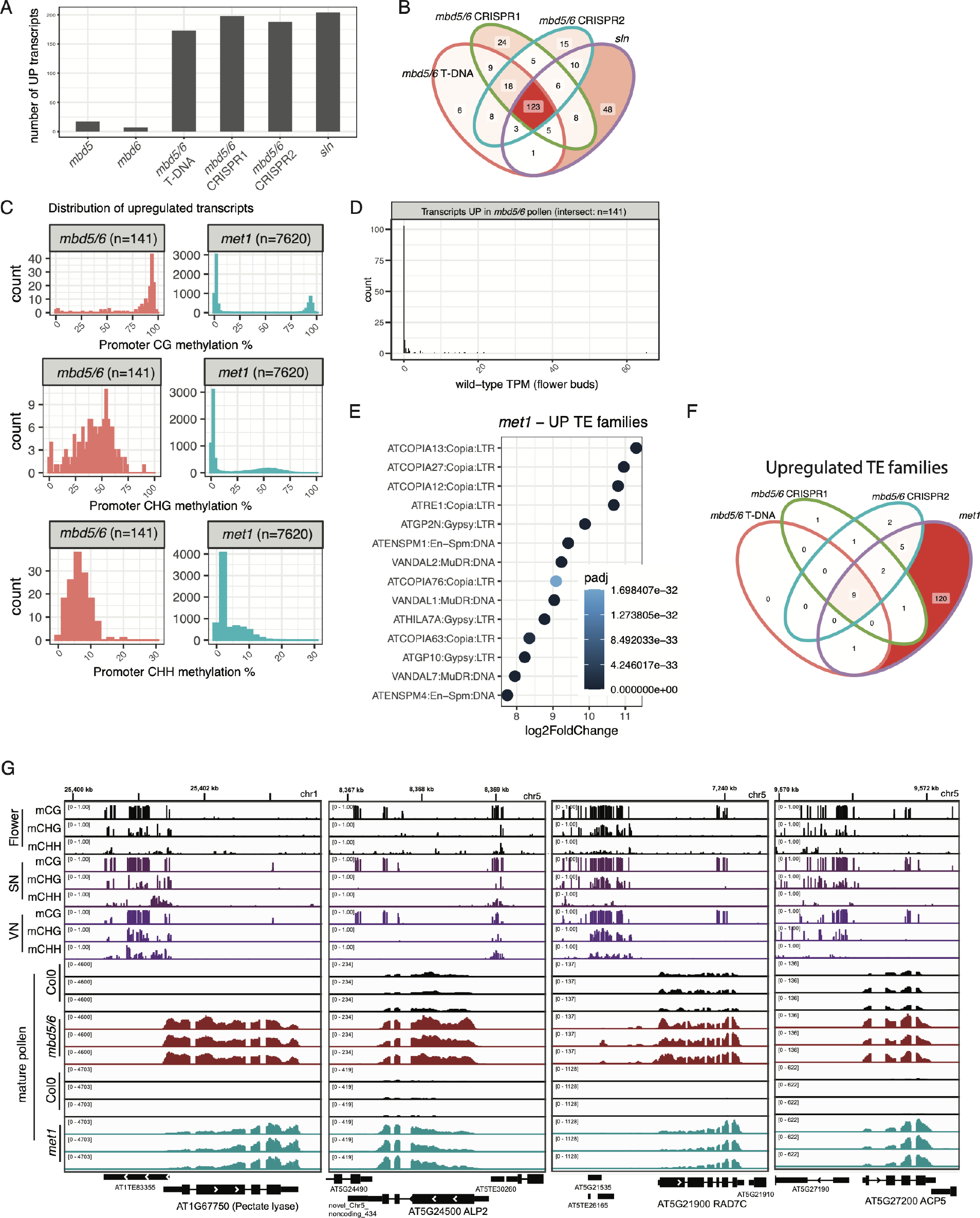
Features of the MBD5/6 targets in mature pollen. A) Number of upregulated transcripts detected by RNA-seq in mature pollen in the indicated mutants. B) Venn diagram showing the overlap between the upregulated transcripts in different mutants. C) Distribution of the promoters that are upregulated either in *mbd5/6* or in *met1* pollen based on their wild-type flower buds methylation levels in CG, CHG, or CHH context. D) Distribution of wild-type expression levels in flower buds for the *mbd5/6* upregulated transcripts. E) Analysis of upregulated TE families in *met1* (mature pollen). Only the top 14 TE families are displayed. F) Venn diagrams of overlaps between the mature pollen upregulated TE families in different mutants. G) Genome browser tracks of mature pollen RNA-seq in *mbd5/6* T-DNA and *met1-3*. Wild-type methylation tracks are shown as reference: flower data is from (Ichino et al., 2021), vegetative nucleus (VN) and sperm nucleus (SN) data is from (Ibarra et al., 2012).

**Figure S5 – related to Figure 3.**
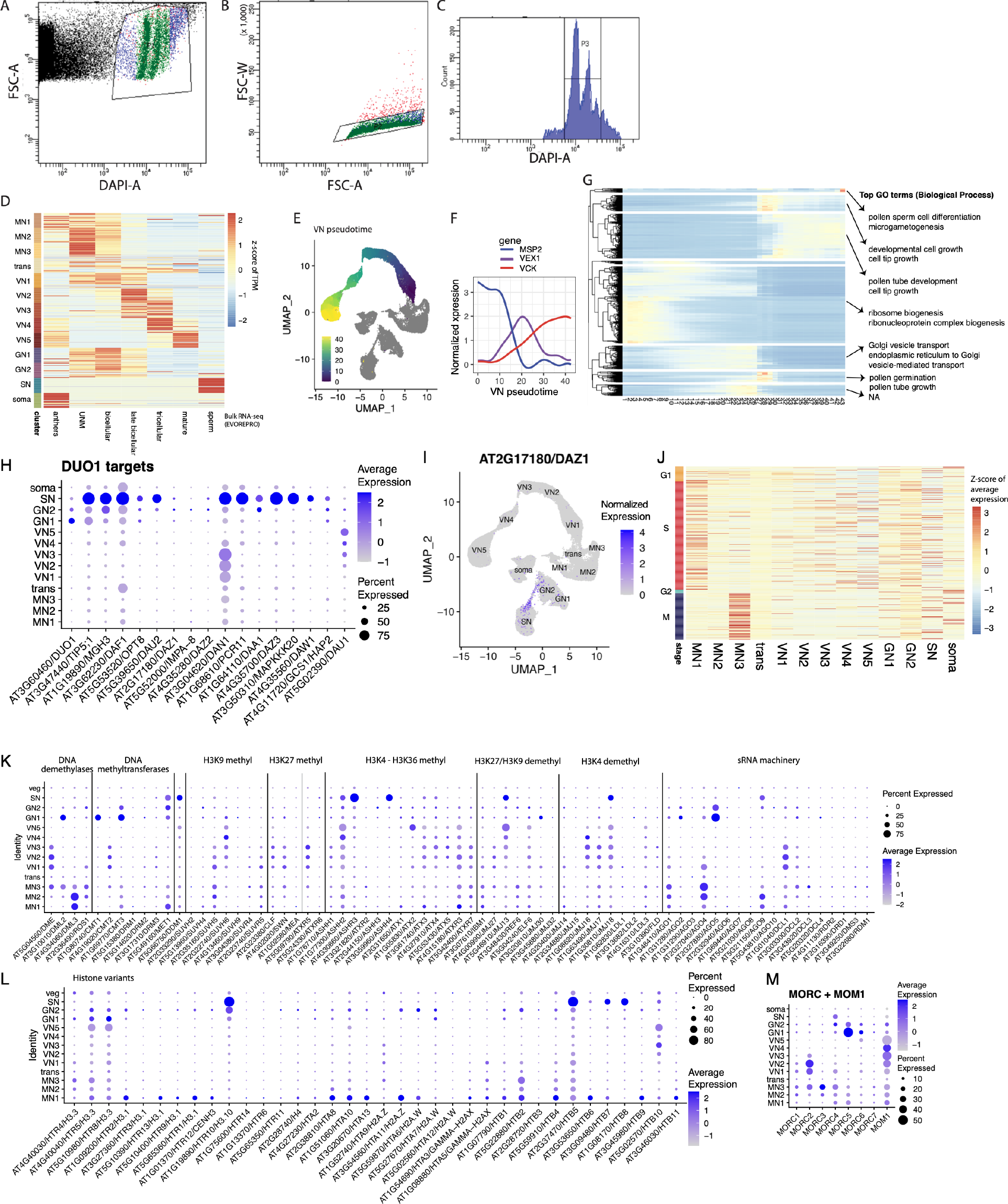
Characterization of snRNA-seq clusters. A,B,C) Representative plots showing the parameters used to isolate and purify the pollen nuclei for snRNA-seq. The gates in panel A were used to separate the DAPI positive nuclei from the small debris, the gates in panel B were used to remove nuclei aggregates (red). The P3 population (Panel C) was sorted in a single tube and used for sn-RNA-seq. The two peaks likely correspond to the large and small nuclei (VN and SN/GN) as the different chromatin structure can lead to a different staining intensities. D) Heatmap of Col0 bulk RNA-seq datasets downloaded from https://evorepro.sbs.ntu.edu.sg/ (Julca et al., 2021). Rows correspond to the top 20 markers for each cluster (ranked by average log2FC) of the snRNA-seq data from this study (the list of markers is available in Table S3). E) UMAP displaying all the wild-type nuclei with their assigned pseudotimes in the VN developmental trajectory (color code). F) Expression pattern along the VN pseudotime for an early VN (*MSP2*), a mid VN (*VEX1*) and a late VN (*VCK*) gene. The lines represent a smoothed trend (ggplot *geom_smooth*) of the log-normalized expression of each gene. G) Heatmap showing 3,135 genes that change as a function of pseudotime (see Methods). The genes were split into 8 groups based on hierarchical clustering. The top ranked GO term for each group is shown (full list in Table S5). H) Dotplot of cluster specific expression profiles for DUO1 and a list of genes that were previously annotated as DUO1 targets (Borg et al., 2011). The dot size represents the percentage of cells in which the gene was detected, the dot color represents the scaled average expression for each cluster. I) UMAP showing the expression pattern of DAZ1, which is specifically expressed in the transition between GN2 and SN. J) Heatmap showing the scaled average expression pattern across all snRNA-seq clusters for a list of previously published cell-cycle marker genes (Menges et al., 2003). Only the genes with a standard deviation higher than 0.2 across all clusters were displayed (n=405). The MN1 and MN3 clusters show a clear enrichment for S-phase and M-phase markers respectively. K,L) Dotplot of cluster specific expression profiles for the indicated groups of genes (see panel H).

**Figure S6 – related to Figure 4.**
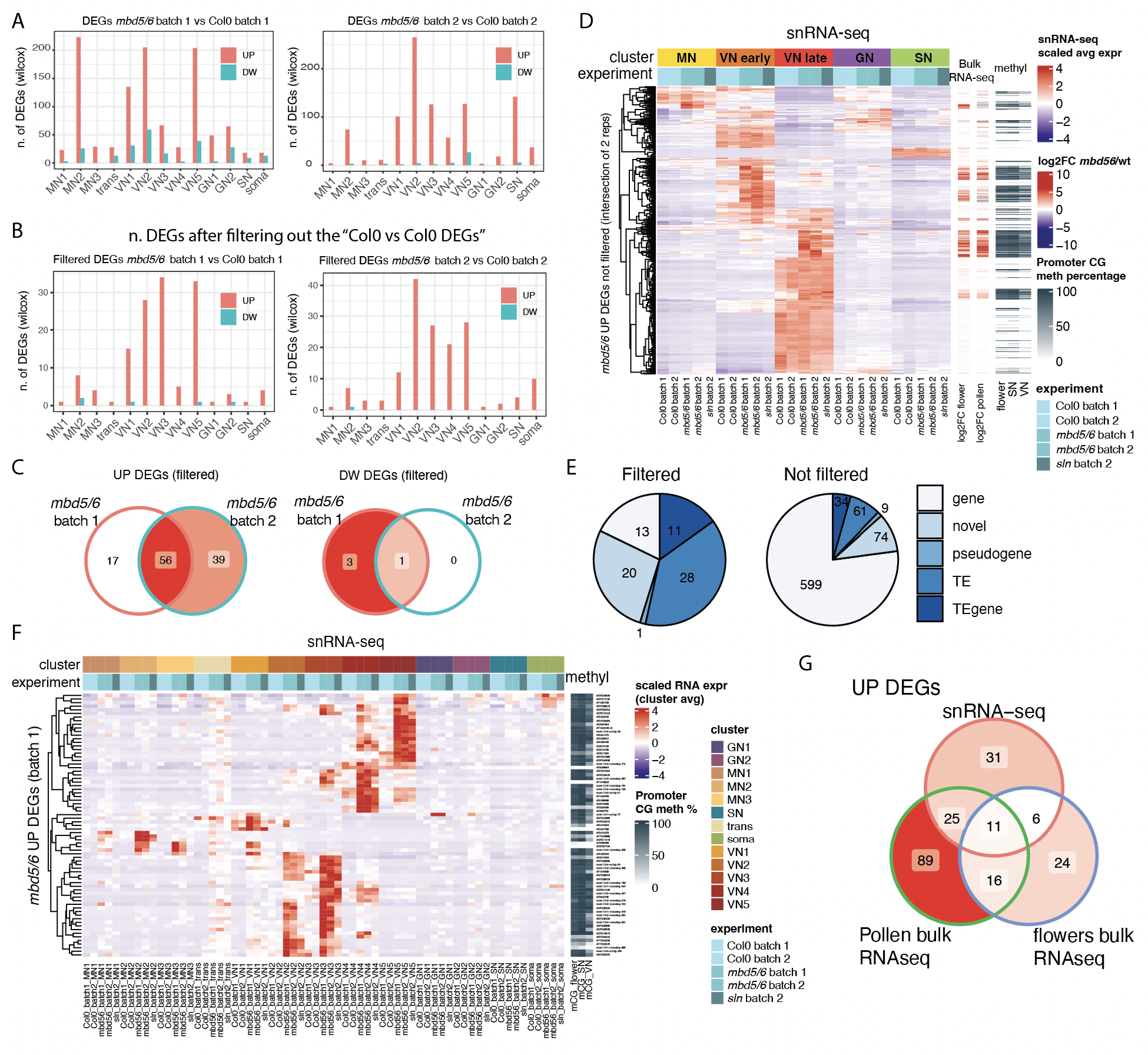
Blacklisting the high variability genes allows detection of high-confidence DEGs with snRNA-seq data. A,B) Barplots of the *mbd5/6* DEGs obtained without filtering (A) or after filtering out the DEGs obtained when comparing Col0 samples to each other (B) (see Methods for details). C) Overlaps of upregulated or downregulated DEGs in two independent experiments. D) Heatmap representation of the not-filtered *mbd5/6* upregulated genes (intersection between batch 1 and batch 2). Shown is the snRNA-seq scaled expression level of the cluster averages in the indicated samples. For each gene, the log2 fold-change obtained by bulk RNA-seq in flower buds or pollen is shown on the right. The last three columns on the right indicate the wild-type CG methylation percentage at the promoters of each gene (600 bp windows centered at the TSS). The VN and SN BS-seq data is from (Ibarra et al., 2012). The genes with a strong positive fold-change in bulk RNA-seq tend to be promoter methylated. E) Classification of the *mbd5/6* upregulated DEGs obtained by snRNA-seq (union of all clusters) either with or without filtering. The “novel” genes were identified via a transcript reannotation based on all the mature pollen RNA-seq datasets included in this study (see Methods and Figure S3). F) Heatmap representation of the union of the *mbd5/6* upregulated genes obtained in each cluster (filtered). Shown is the snRNA-seq scaled expression level of the cluster averages in the indicated samples. The methylation data is the same as in panel D. G) Venn diagram of overlaps between snRNA-seq batch 1 and bulk-RNAseq upregulated DEGs.

**Figure S7– related to Figure 5:**
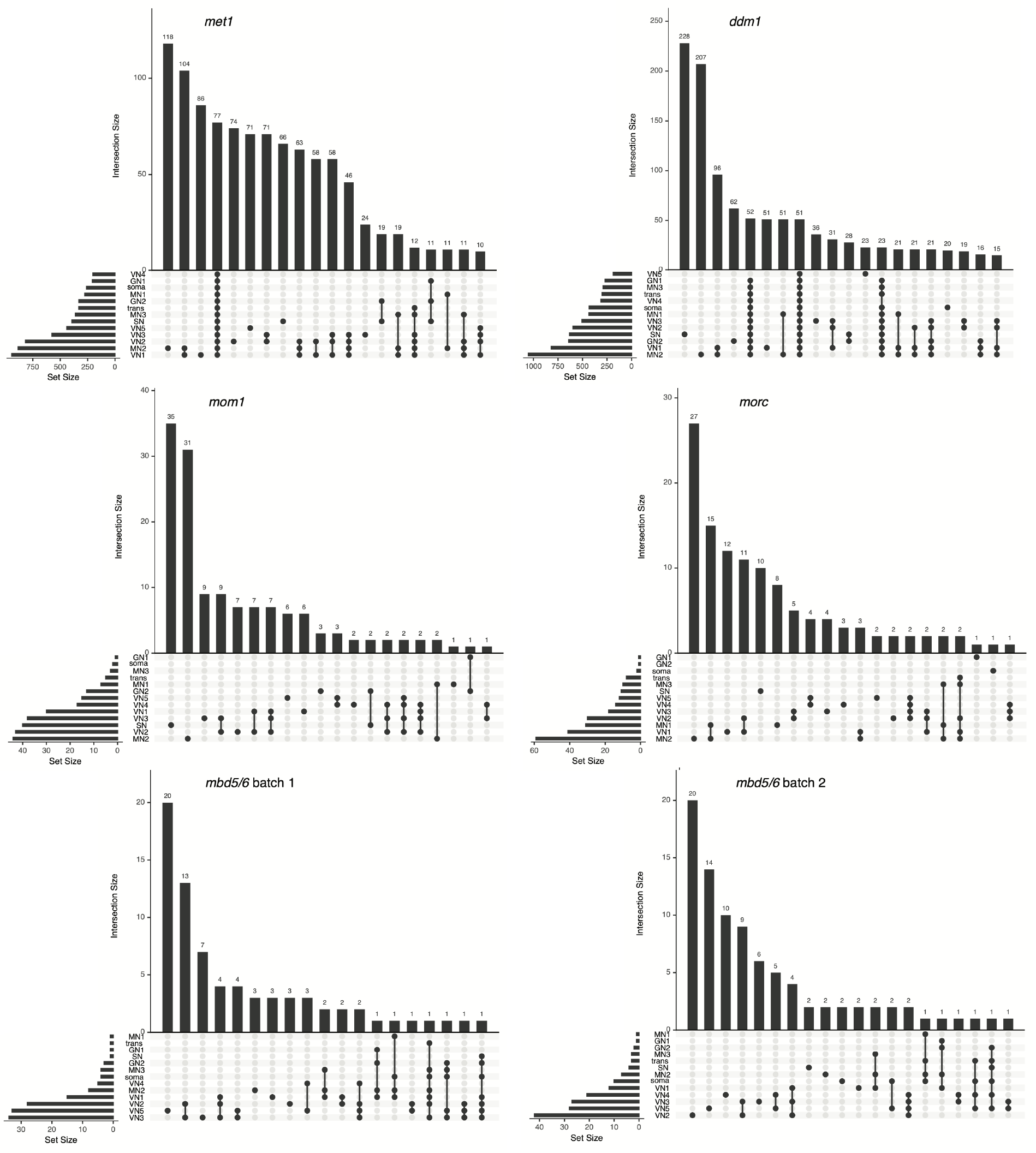
Differential gene expression analysis at individual nuclei clusters. Upset plots showing the intersection of the upregulated transcripts obtained in each nucleus type, for the indicated genotypes.

